# A shared slowing axis and phenotype-specific rhythms link clinical EEG to FDG-PET dementia phenotypes

**DOI:** 10.64898/2026.11.09.764237

**Authors:** Nihal Satyadev, Venkatsampath Gogineni, Nick Corriveau-Lecavalier, Gemeng Zhang, Benjamin Brinkmann, Yogatheesan Varatharajah, Gregory S. Day, Jennifer L Whitwell, Keith A. Josephs, Kejal Kantarci, Bradley F. Boeve, Ronald C. Petersen, Michael Breakspear, Val J. Lowe, David T. Jones

## Abstract

Early and accurate identification of neurodegenerative disorders has notable implications for treatment and quality of life. Whereas neuroimaging-based biomarkers can accurately identify neurodegenerative subtypes, EEG represents a low-cost alternative that could expand access to dementia care. Here, we investigated whether six data-driven EEG latent factors, derived by nonnegative tensor decomposition of clinical EEGs, differ systematically across FDG-PET-defined neurodegenerative phenotypes.

In 455 patients with EEG and FDG-PET acquired within one year of each other, factor expression was compared across 14 diagnostic classes assigned by StateViewer, a validated artificial intelligence framework for FDG-PET interpretation. To separate disease-general severity from phenotype-specific topography, each factor was also regressed on a subject-level slowing index. Finally, we related the factors to regional FDG uptake and to neuropsychological performance (n = 98) and assessed their potential utility for single-subject classification.

Four factors reflecting neurophysiological oscillations (posterior alpha, Rolandic beta, frontotemporal delta–theta and posterior theta–slow alpha) varied significantly across phenotypes, with attenuation of faster and augmentation of slower rhythms most pronounced in dementia with Lewy bodies (DLB), dysexecutive Alzheimer disease (AD) and posterior cortical atrophy, whereas factors dominated by eye-movement and muscle artifact showed no robust phenotypic association. Alpha and beta effects were largely attributable to overall slowing, whereas posterior theta–slow alpha became more phenotype-specific after adjustment (η² 0.082 to 0.147), dissociating the Alzheimer disease spectrum from frontotemporal syndromes. The slowing index tracked with an FDG-PET pattern highly associated with DLB across the cohort, irrespective of assigned phenotype (r = 0.51, *P* < 0.001). Consistent with a model in which dementia phenotypes reflect altered hippocampal–neocortical oscillatory coupling (stochastic latent oscillatory diffusion), frontotemporal delta–theta tracked a limbocortical axis of metabolic pattern expression, rising with neocortical and falling with limbic-predominant involvement, independent of overall slowing and phenotype. Each factor mapped onto a distinct metabolic topography, with frontotemporal delta–theta resembling Alzheimer-type hypometabolism and posterior theta–slow alpha the medial occipital hypometabolism of DLB. In terms of neuropsychological performance, Rolandic beta was associated with executive functions, and frontotemporal delta–theta with memory, visuospatial and global performance. Using these four factors alone, an exploratory logistic regression classifier separated cognitively unimpaired participants from DLB, posterior cortical atrophy, typical AD, and dysexecutive AD (area under the curve > 0.8) but poorly discriminated frontotemporal dementia phenotypes canonically associated with 4-repeat tau.

Thus, data-driven decomposition of clinical EEG separates oscillatory activity from artifact into factors that track metabolic phenotype, regional metabolism and cognition, capturing both overall slowing and phenotype-specific patterns. This work informs the further development of scalable, biology-informed EEG-based phenotyping of neurodegenerative disease.

## Introduction

Neurodegenerative disorders of cognition and behavior are disorders of large-scale brain functional physiology.^1,2^ Although clinical syndromes such as Alzheimer disease (AD) dementia, dementia with Lewy bodies (DLB), and frontotemporal dementias often present with overlapping cognitive and behavioral features, each arises from distinct biological processes that selectively disrupt distributed cortical and subcortical circuits.^3^ Therefore, improving trans-diagnostic precision requires approaches that move beyond descriptive syndromes toward measures that directly frame functional brain physiology as a common pathological mechanism. Although expert clinical judgment remains the standard, it is inherently subjective and difficult to scale, motivating the development of biologically grounded clinical decision support systems that leverage machine learning methodologies to improve diagnostic quality in neurological practice.^4^

Our group has previously demonstrated the utility of fluorodeoxyglucose positron emission tomography (FDG-PET) as a highly accurate framework that can be incorporated into clinical decision support systems to distinguish between nine different neurodegenerative syndromes.^5^ However, FDG-PET can be resource-intensive and not universally available, limiting use in screening, longitudinal monitoring, or early disease detection in routine care settings. In addition, static measurements of functional physiology are insensitive to important aspects of functional dynamics that are likely relevant for neurodegenerative physiology. Scalp electroencephalography (EEG), by contrast, is inexpensive, widely accessible, and directly reflects neuronal population activity, making it an attractive candidate for scalable neurodegenerative phenotyping related to functional brain physiology. Additionally, EEG has previously been shown to distinguish between various subtypes of AD and related dementias.^6–8^ However, EEG is still limited in its widespread clinical utility as it is currently only referenced as a biomarker of utility in the diagnostic criterion for one neurodegenerative disorder – DLB.^9^ More details are needed to clarify the role of neuronal oscillations in the biological underpinnings of neurodegeneration to expand the clinical applications of EEG.

Neuronal oscillations, or brain rhythms, are categorized by five major bands (delta, δ, 0.1–4 Hz; theta, θ, 4–8 Hz; alpha, α, 8–12 Hz; beta, β, 12–30 Hz; and gamma, γ, >30 Hz) and reflect coordinated timing across neuronal populations that depend on intact synaptic transmission, inhibitory interneuron function, and neuromodulatory input.^10–12^ Disruption of these processes leads to systematic, frequency-specific alterations in large-scale network dynamics that are detectable using EEG. The relationship between dementia and changes in brain rhythms can be conceptualized in emerging oscillatory computational models of hippocampal-neocortical interaction (i.e., stochastic latent oscillatory diffusion, SLOD).^13^ Across neurodegenerative disorders, prior studies have consistently demonstrated shifts in resting-state oscillations, including attenuation of faster rhythms and relative increases in slower activity.^14,15^ Crucially, oscillatory abnormalities are not uniform across neurodegenerative syndromes or even across phenotypes within a single diagnostic category. Hence, analyses based on predefined frequency bands, as often done by traditional quantitative EEG methods, offer a fragmented view of brain activity confined to individual spectral ranges, limiting the detection of EEG patterns that extend across multiple frequencies. In addition, clinical EEG recordings often contain non-neuronal artifacts that can confound measures of power and synchrony.

Data-driven modeling approaches provide a principled method of addressing these limitations. By decomposing EEG signals without predefined features, latent factor models can identify population-level spectral and spatial patterns that reflect fundamental sources of electrophysiological variance.^16,17^ Importantly, this framework allows for explicit separation of oscillatory components associated with neuronal biology from components dominated by non-neuronal artifact, enabling direct evaluation of which EEG features carry disease-relevant information.

Here we investigate whether data-driven latent factors in resting, eyes-closed EEG differ systematically across neurodegenerative phenotypes defined by FDG-PET, as classified by a clinically validated artificial intelligence model grounded in a global functional state space formulation of brain function.^1,5^ Using resting-state EEG from a large cohort of patients with paired EEG and FDG-PET data, we derive and examine the association between six population-level EEG factors and fourteen FDG-PET phenotypic classifications. We hypothesized that neurophysiologic oscillatory factors would demonstrate phenotype-specific variation, while factors dominated by non-neuronal artifact would not, and that distinct neurodegenerative metabolic phenotypes exhibit distinct electrophysiological signatures. By anchoring EEG neurophysiologic features to FDG-PET–defined biology, this work aims to expand the physiologic associations of EEG oscillations in neurodegenerative disease and to establish a foundation for scalable, biology-informed EEG-based phenotyping with further model development.

## Methods

### Participants

Participants for this study were drawn from four sources within the Mayo Clinic Enterprise: the Mayo Clinic Alzheimer’s Disease Research Center, the Mayo Clinic Study of Aging, the Neurodegenerative Research Group, and patients from the neurology clinic who authorized use of their health records for research but were not part of any specific study. Data usage from these participants was approved by the Mayo Clinic Institutional Review Board. A medical record search was performed for patients over the age of 18 who had both a routine EEG and FDG-PET done within one year, which resulted in 1,073 pairs. If multiple studies were done, only the EEG/FDG-PET combination with the closest temporal relationship were selected, yielding 875 pairs. Only high confidence FDG-PET class selections (i.e., StateViewer class diagnostic assignments with *P* < 0.01) were retained, resulting in a final cohort of 455 paired EEG/FDG-PET studies. Restricting the analysis to high-confidence assignments minimizes misclassification of phenotype labels, so that between-group differences in EEG factor expression reflect the underlying neurodegenerative pattern rather than diagnostic noise from ambiguous or mixed metabolic profiles. To assess the effect of this filtering, primary analyses were repeated in the full cohort of the 875 paired studies.

### FDG-PET Imaging

FDG-PET imaging was performed using a PET/CT scanner (GE Healthcare) following a 30-minute uptake period during which participants rested in a dimly lit room. Image acquisition lasted 8 minutes and consisted of four dynamic frames (2 minutes each), preceded by a low-dose CT transmission scan. FDG-PET images were processed using an MRI-independent pipeline.^5^ This included non-linear symmetric diffeomorphic registration to the Mayo Clinic adult lifespan template, normalization of uptake values to the pons to generate standardized uptake value ratio images, and spatial smoothing with a 6 mm full width at half maximum Gaussian kernel.

### FDG-PET Stateviewer

As previously described,^5^ 3,671 FDG-PET images from Mayo Clinic (Rochester, Minnesota) were used to create StateViewer, an imaging-based clinical decision support system. StateViewer treats dementia diagnoses as a multiclass, multilabel problem, reflecting the potential for overlapping neurodegenerative pathologies. The model employs a k-nearest neighbor (k-NN) approach in which each query scan is compared against a reference population with expert-assigned clinical phenotype labels. For each disease class, enrichment within the local k-NN neighborhood is quantified using odds ratios and one-sided Fisher exact test p-values, allowing multiple phenotype detections per individual while accounting for class imbalance.

Model outputs consist of zero or more detected phenotypes ranked by statistical evidence, enabling dynamic adjustment of sensitivity and precision via p-value thresholds. Model performance was evaluated using sensitivity, precision, and discovery rate metrics optimized for clinical decision support, with interpretability provided through visualization of nearest-neighbor relationships and representative FDG-PET metabolic patterns derived from population-level feature space. Fourteen phenotypes are currently classified in the StateViewer framework: AD dementia,^18^ behavioral variant frontotemporal dementia,^19^ corticobasal syndrome,^20^ cognitively unimpaired (CU), dysexecutive AD,^21^ DLB,^9^ limbic-predominant amnestic neurodegenerative syndrome (LANS),^22^ normal pressure hydrocephalus,^23^ posterior cortical atrophy (PCA),^24^ primary progressive apraxia of speech,^25^ progressive supranuclear palsy,^26^ logopenic variant primary progressive aphasia, nonfluent variant primary progressive aphasia, and semantic dementia.^27^

### EEG Analysis

EEGs were recorded using the XLTEK EMU40EX headbox (Natus Medical Inc., Oakville, Ontario, Canada) according to the standard 10–20 localization system^28^ at a sampling rate of 256 Hz. As previously described,^17^ 12,176 clinical EEG recordings of 11,001 adult patients (≥18 years old) who underwent routine clinical EEG study at the Mayo Clinic (Rochester, Minnesota) between the years 2011 and 2021 were used to create latent factors in an unsupervised fashion. An automated algorithm selected between four and six 10-second-long epochs in the eyes-closed awake state. The standard 19 EEG channels were maintained for analysis. All channels underwent Welch fast Fourier transform approach to estimate the power spectral density. These matrices were stacked to form a 3D tensor of dimension 12,176 x 19 x 45. Nonnegative parallel factor analysis (i.e., canonical polyadic) decomposition yielded six factors, each consisting of 3 rank-1 tensors corresponding to EEG record, channel, and frequency dimensions. Factors one through six were previously hypothesized to correlate with eye movement artifact, posterior alpha rhythm, Rolandic beta activity, frontal/temporal delta and slow theta rhythm, bitemporal muscle artifact, and posterior theta and slow alpha rhythm, respectively.^17^

### FDG-PET Metabolic Pattern Expression

Metabolic pattern expression (MPE) values were derived from representative images corresponding to each diagnostic class from StateViewer.^5^ For each class, a median vector was computed in principal component space as the minimizer of total cosine distance, obtained operationally by normalizing constituent vectors to unit length and taking their mean direction. In this manner, metabolic pattern expression represents a whole-brain pattern aligned or anti-aligned with membership in that diagnostic class. MPE vectors were applied at the participant level across all cohorts to extract class-specific expression values. Because posterior EEG slowing is a recognized supportive biomarker of DLB,^9^ we hypothesized a priori that EEG slowing would track DLB MPE specifically.

### Between-phenotype Comparisons

Levels of each of the six EEG-derived latent factors were compared across the 14 FDG-PET–derived phenotypic classes using one-way analysis of variance (ANOVA). When the omnibus ANOVA was significant, pairwise post-hoc comparisons were conducted using two-sided t-tests. To control the family-wise error rate across the 91 pairwise comparisons within each factor, p-values were adjusted using the Holm step-down procedure. Homogeneity of variance was evaluated using Levene’s test, and Welch ANOVA was used in cases where homogeneity assumption was violated.

Factor loadings were standardized against the CU class (z = (x − μ_CU)/σ_CU). A slowing index was computed per subject as the sign-oriented mean of the four neurophysiologic factors: (−z_F2 − z_F3 + z_F4 + z_F6)/4. Factor 2 and factor 3 were reversed given that they are faster rhythms, and artifact factors (1 and 5) were excluded. To distinguish phenotype-specific topography from disease-general slowing, each factor was regressed on this index across the cohort and residuals averaged within phenotype. Residual deviations were tested against a family-wise threshold derived from 5,000 label permutations using a max-statistic null across all factor-by-phenotype cells.

Finally, to determine whether EEG tracks metabolic phenotype expression continuously rather than only across diagnostic categories, DLB MPE was regressed on the slowing index by ordinary least squares across all subjects, irrespective of assigned phenotype label. To assess specificity, the association between the slowing index and each remaining class-specific MPE was then tested in a model that also included DLB MPE. To assess the SLOD framework hypothesis^13^ that phenotypes reflect differences in cortical-hippocampal interaction and the associated oscillatory dynamics, we derived a limbocortical axis (LCA) by subtracting LANS MPE from dysexecutive AD MPE and associated each non-artifactual factor with this index. In this paper, statistical significance was set at α = 0.05 throughout and all analyses were conducted in Python (version 3.14).

### Voxelwise Analysis and Decoding

To characterize each factor’s metabolic correlates without reference to predefined templates, voxelwise regression was performed in which FDG-PET uptake at each voxel was modeled as a function of the four neurophysiologic EEG factors (factors 2, 3, 4, and 6), entered simultaneously in a single multivariable model across all subjects in the primary cohort (n = 455) with age and sex included as covariates. Because the factors were entered jointly, each resulting β map reflects the association between regional metabolism and one factor while holding the other three constant. A subsequent meta-analytical decoding was performed for each of the four β maps using the Neurosynth database (www.neurosynth.org).^29^ Consistent with prior efforts,^1^ we used a predefined set of topic-term families (v5-topics-50) and limited analysis to topics related to cognition.^30^ Each of these topics were subsequently associated with each β map and the five most positively and the five most negatively associated topics were presented. Interpretation of these findings are meant to be guided by directionality and relative strength rather than absolute values. As this analysis was done purely for descriptive purposes, no significance values are reported.

By contrast, alignment of each β map with macroscale cortical organization was tested inferentially. Maps were projected to the cortical surface, averaged within 400 cortical parcels^31^ and correlated (Spearman’s ρ) with the principal (G1; unimodal to transmodal) and second (G2; somatomotor to visual) functional connectivity gradients.^32^ Significance was assessed against 10,000 spin tests^33^ and Holm-corrected across the four factors within each gradient (Supplementary Methods).

### Neuropsychology

A subset of patients (n = 98) completed neuropsychological testing within a year of FDG-PET and had results available for review. Scores were drawn from 23 tests. Each score was expressed as a percentage of the maximum possible score, with timed measures inverted so that higher values always indicate superior performance. Tests were assigned to four cognitive domains (1) Memory: Auditory Verbal Learning Test (trial 1, trials 1-5, list B, trial 6, delayed recall); Wechsler Memory Scale logical memory I/II and visual reproduction I/II; (2) Executive Function: Trail Making A/B; Stroop word, color and color–word; Wechsler Adult Intelligence Scale digit span and coding; (3) Language: Boston Naming Test; Controlled Oral Word Association Test; Category fluency; (4) Visuospatial function: Block Design, Judgment of Line Orientation, Picture Completion, Rey–Osterrieth figure. Participants completed different subsets of the battery. To account for this, each test was z-scored within the neuropsychological subset, and a domain composite was calculated as the mean z-score of the available tests in that domain. A global composite was the mean of the available domain composites. Associations between each factor and each composite were estimated with Spearman rank correlation under three models: unadjusted; partial correlation adjusted for age and years of education; and additionally adjusted for StateViewer phenotype (14-level fixed effect), which estimates the within-phenotype association. Partial correlations were obtained by correlating residuals of the rank-transformed variables after regression on the rank-transformed covariates, with degrees of freedom of n − 2 − k. P-values were Holm-corrected within each model across the 20 composite × factor comparisons. Test-level associations were estimated using model 2 (age and education adjusted), with Holm correction across all 92 test × factor comparisons.

### Single-subject Classification

As an exploratory effort, we tested whether the four neurophysiological EEG factors alone could discriminate clinical phenotypes. We defined five classification analyses a priori using the full cohort (*n* = 455). (1) CU (*n* = 50) versus each of the 13 other phenotypes, as 13 separate binary models. (2) CU versus all cognitively impaired participants pooled (*n* = 405). (3) CU versus AD-associated phenotypes (PCA, logopenic variant primary progressive aphasia, typical AD and dysexecutive AD; *n* = 131). (4) CU versus 4 repeat (4R)-associated phenotypes (progressive supranuclear palsy, corticobasal syndrome and primary progressive apraxia of speech, *n* = 64). (5) A four-class model separating AD-associated (*n* = 131), 4R-associated (*n* = 64), DLB (*n* = 47) and semantic dementia (*n* = 37) participants. Multi-class groups were segmented by proteinopathies. AD-associated phenotypes include those associated with amyloid beta plaques and mixed 3R/4R tau intraneuronal neurofibrillary tangles.^34^ 4R-phenotypes are associated with 4R tau with dominant accumulation in glial cells.^35,36^ Semantic dementia has strong neuropathological correlation with TDP-43 Type C pathology, and finally DLB is an alpha-synucleinopathy.^36^ Behavioral variant frontotemporal dementia, nonfluent variant primary progressive aphasia, and LANS were excluded from this analysis due to their heterogeneous association with proteinopathies.^22,37,38^ Normal pressure hydrocephalus was excluded as it is a CSF-dynamics disorder of indeterminate, multifactorial etiology.^39^ To test whether conclusions depended on the learning algorithm, we compared five widely used classifiers: Gaussian naïve Bayes (as used in our prior work),^17^ L2-regularized logistic regression, support vector machine, random forest and gradient boosting. Performance was estimated by stratified five-fold cross-validation, repeated 10 times with different random fold assignments. Each participant’s out-of-fold scores were averaged across repeats, and from these we computed the area under the receiver operating characteristic curve (AUC) as the primary metric. We also computed balanced accuracy, sensitivity and specificity at the class-balanced threshold. P-values were obtained by label-permutation testing of the primary (logistic regression) model. They were corrected for multiple comparisons with the Holm–Bonferroni procedure, within analysis (1) (13 tests) and across analyses (2)–(5) (4 tests). More details on the machine-learning methodology are presented in the Supplementary Methods.

## Results

A total of 455 patients with paired routine EEG and FDG-PET performed within one year were included in the analysis (Fig. 1; Table 1). The median age at symptom onset was 62.0 years (range 18.2–90.4), with a near-equal distribution of sex (female n=220, 48.4%). The cohort was predominantly non-Hispanic White (n=414, 91.0%). The median interval between EEG and FDG-PET was 2 days (IQR 1-22). Normal pressure hydrocephalus comprised the largest group (n=67, 14.7%), followed by CU (n=50, 11.0%), and dysexecutive AD (n=48, 10.5%). Table 1 lists the complete set of classes.

**Figure 1:**
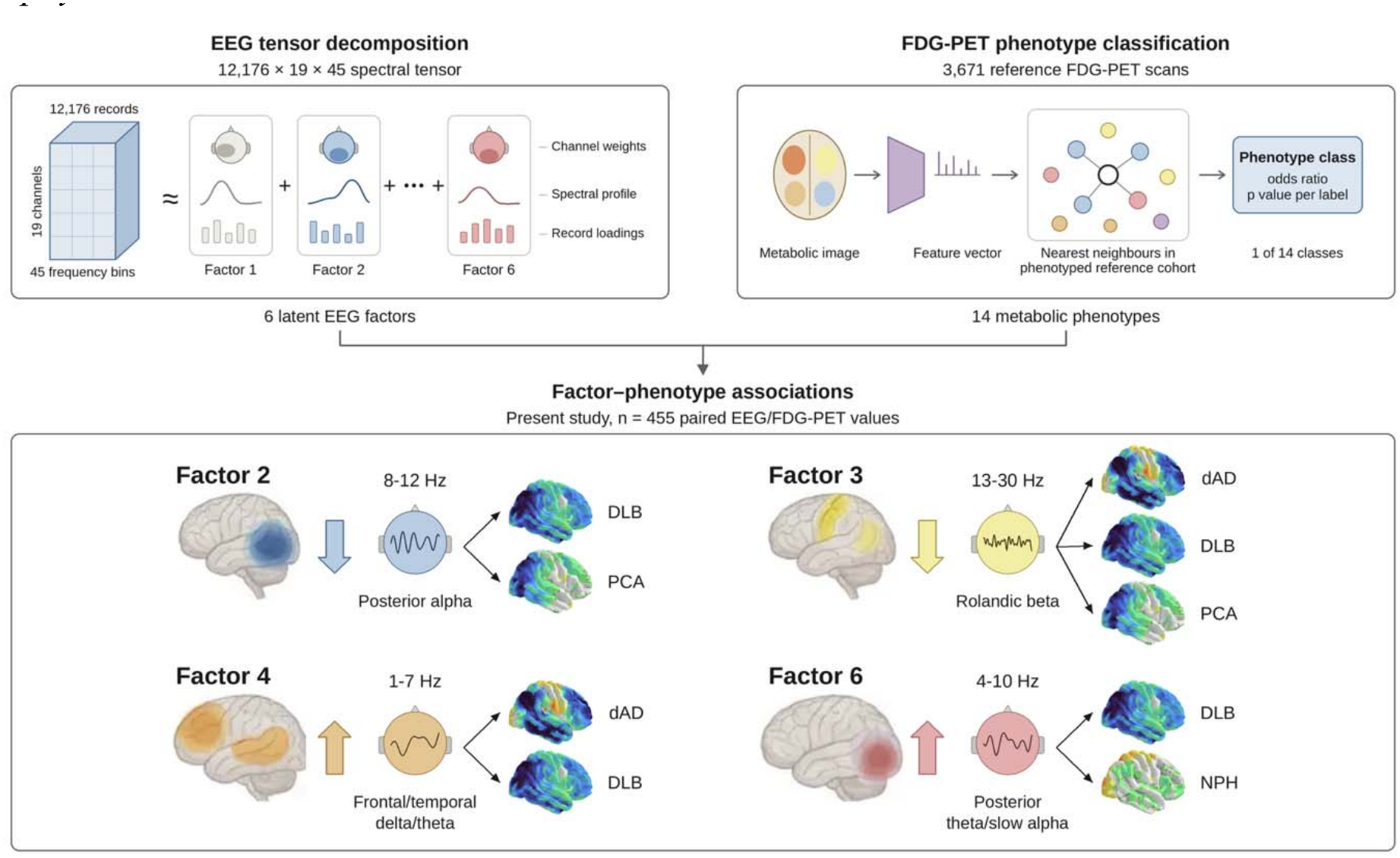
Workflow linking data-driven EEG oscillatory factors to FDG-PET metabolic phenotypes. **Top:** the two independently derived inputs: PARAFAC decomposition of the clinical EEG spectral tensor (left)^17^ and StateViewer FDG-PET phenotype classification (right)^5^ which contribute the six latent factors and the 14 phenotype classes, respectively. **Bottom:** FDG-PET phenotypes and EEG are linked for the first time in the present cohort. The four neurophysiologic factors presented with the phenotypes each is associated with. Lateral brain and head schematics indicate hypothesized topography and frequency band. Vertical arrows give the direction of the factor loading in the associated phenotypes relative to the cognitively unimpaired class. Cortical surface renders show a single-subject example FDG-PET expression for each phenotype (left lateral view; *z* scale −3.5 to +3.5). *dAD, dysexecutive Alzheimer disease; DLB, dementia with Lewy bodies; NPH, normal pressure hydrocephalus; PCA, posterior cortical atrophy*.

FDG-PET phenotypic class accounted for 12.5% of the variance in latent factor 2 (F(13, 441) = 4.835, *P* < 0.001, η² = 0.125, Fig. 2A). Post-hoc pairwise comparisons with Holm correction identified significant differences between the DLB class versus the normal pressure hydrocephalus (t(441) = −4.58, adjusted *P* = 0.001) and CU (t(441) = −5.40, adjusted *P* < 0.001) classes. Differences were also observed between the PCA class versus the normal pressure hydrocephalus (t(441) = −4.94, adjusted p < 0.001), progressive supranuclear palsy (t(441) = − 3.84, adjusted *P* = 0.024), behavioral variant frontotemporal dementia (t(441) = −3.74, adjusted *P* = 0.035), and CU (t(441) = −5.67, adjusted *P* < 0.001) classes, and between the dysexecutive AD and CU classes (t(441) = −3.92, adjusted *P* = 0.015). Phenotype also accounted for a significant portion of variance in EEG latent factor 3 (F(13, 132.95) = 5.945, *P* < 0.001, η² = 0.154), factor 4 (F(13, 441) = 9.324, *P* < 0.001, η² = 0.216), and factor 6 (F(13, 133.07) = 4.711, *P* < 0.001, η² = 0.082). The full pattern of pairwise comparisons is summarized in the corresponding matrix (Fig. 2). Phenotype did not explain significant variance in factor 1 (F(13, 441) = 1.37, *P* = 0.169, η² = 0.039) and was nominally significant for factor 5 (F(13, 441) = 1.91, *P* = 0.027, η² = 0.053) but not after Holm correction across the six factors (*P* = 0.054); no pairwise comparison survived correction for either factor (Supplementary Fig. 1). Filtering FDG-PET classes to only those with high-confidence (*P* < 0.01) resulted in largest dropouts in those with primary progressive apraxia of speech, corticobasal syndrome, and LANS (Supplementary Table 1). However, the main factor differences held directionally in the full set of 875 cases (Supplementary Fig. 2).

**Figure 2:**
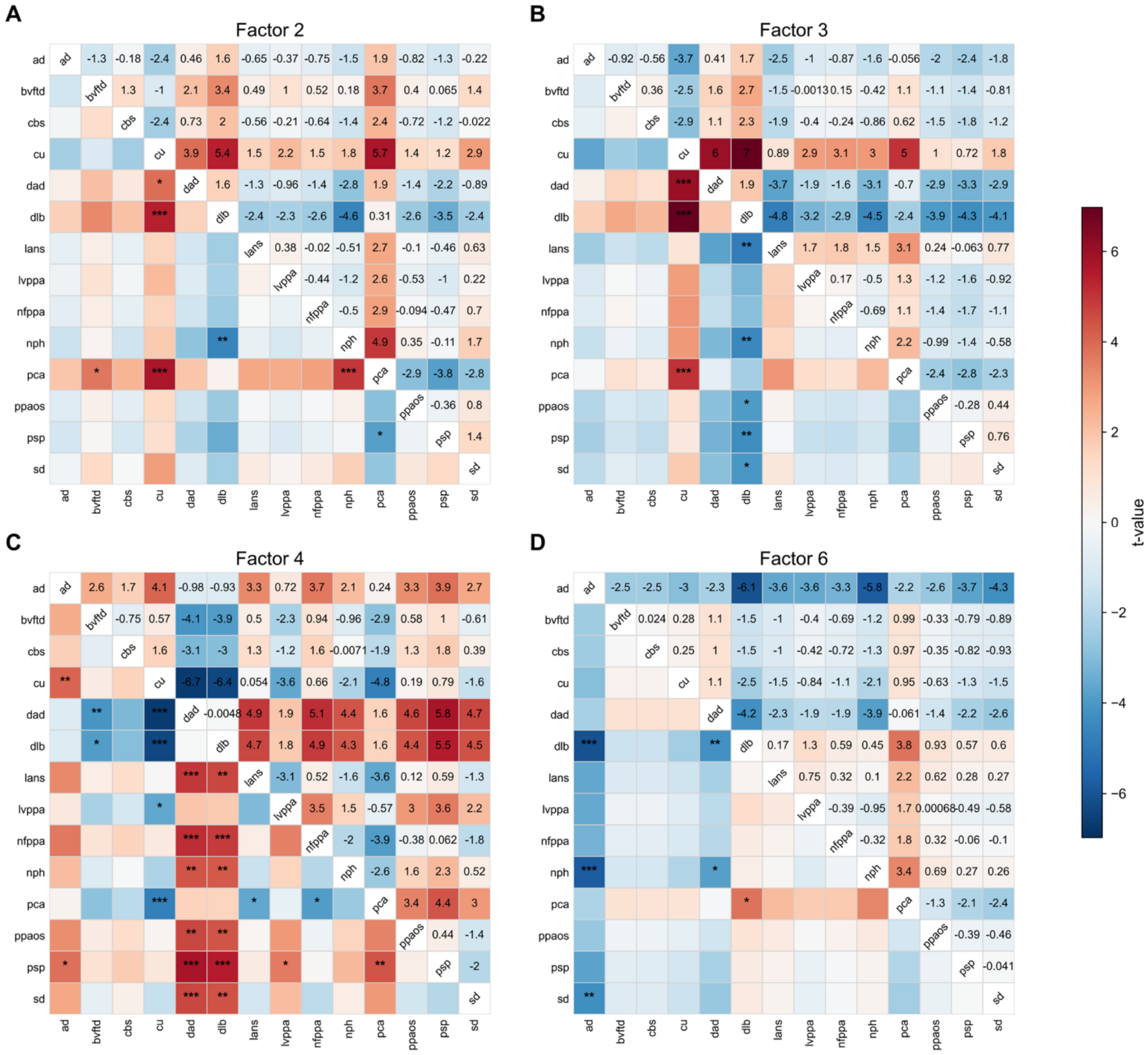
Pairwise differences in latent EEG factor scores across clinical syndromes. Each panel shows all pairwise comparisons of subject-level scores for one latent EEG factor among the 14 clinical groups: **(A)** Factor 2, **(B)** Factor 3, **(C)** Factor 4, and **(D)** Factor 6. Comparisons are presented as 14×14 matrices, with the upper triangle displaying signed t statistics (row minus column comparisons) and the lower triangle indicating statistical significance based on Holm-adjusted p-values (*p<0.05, **p<0.01, ***p<0.001). Cell shading encodes the t statistic (red, higher scores in the row group; blue, lower). For clarity of presentation, Factors 1 and 5, for which no pairwise comparison remained significant after Holm correction, are reported in Supplementary Figure 1. *AD, Alzheimer disease; bvFTD, behavioral variant frontotemporal dementia; CBS, corticobasal syndrome; CU, cognitively unimpaired; dAD, dysexecutive Alzheimer disease; DLB, dementia with Lewy bodies; LANS, limbic-predominant amnestic neurodegenerative syndrome; lvPPA, logopenic variant primary progressive aphasia; nfvPPA, nonfluent variant primary progressive aphasia; NPH, normal pressure hydrocephalus; PCA, posterior cortical atrophy; PPAOS, primary progressive apraxia of speech; PSP, progressive supranuclear palsy; SD, semantic dementia*.

Expressing factor levels relative to the CU class placed all fourteen phenotypes on a common severity axis (Fig. 3A). Each disease phenotype showed a positive slowing index, ranging from +0.15 in progressive supranuclear palsy to +1.10 in DLB, with PCA (+0.74) and dysexecutive AD (+0.79) also at the slower end. DLB showed the most extreme profile on every neurophysiologic factor (factor 2: −1.01, factor 3: −1.39, factor 4: +1.55, factor 6: +0.44 standard deviations). Per-subject distributions confirmed that this ordering reflects genuine group separation rather than outlying individuals (Fig. 3C). Because much of this structure reflects a disease-general shift toward slower activity, each factor was regressed on the subject-level slowing index and the residuals re-examined (Fig. 3B). Variance attributable to phenotype decreased for factor 2 (η² 0.125 → 0.047), factor 3 (0.154 → 0.035), and factor 4 (0.216 → 0.118), while it increased for factor 6 (0.082 → 0.147). AD, PCA, and dysexecutive AD demonstrated increased factor 4 and reduced factor 6 after severity adjustment. In contrast, 4R tau syndromes and behavioral variant frontotemporal dementia broadly demonstrated inverse values among those two factors. DLB showed near-zero residuals on all four neurophysiologic factors (|residual| ≤ 0.15).

**Figure 3:**
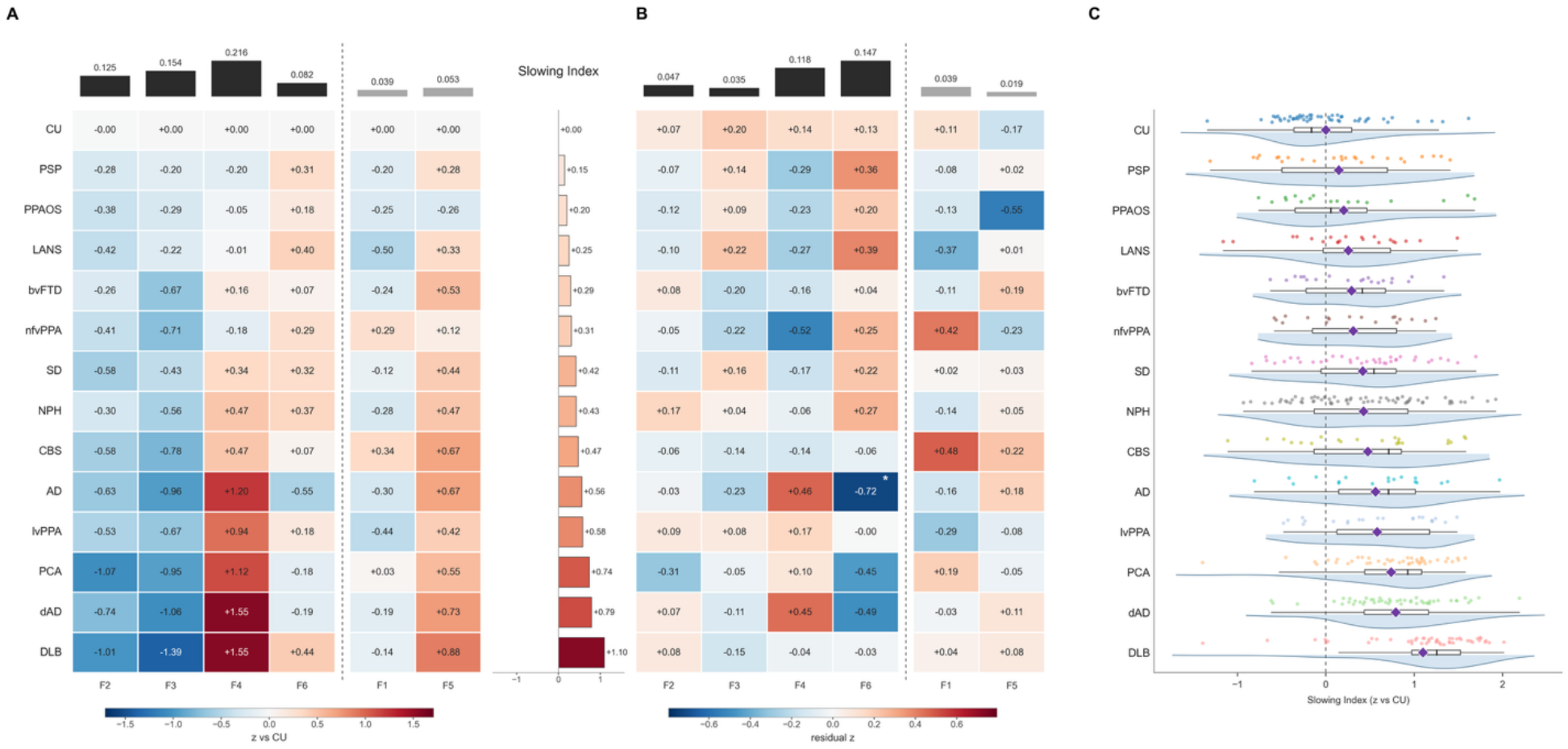
EEG latent factor expression across FDG-PET–defined metabolic phenotypes. **(A)** Mean factor level for each phenotype, expressed in standard deviation units relative to the CU class. Arrows in the column headers indicate the pathological direction of each factor. Bars above each column give η², the proportion of variance in that factor attributable to phenotype. **(Center)** Group mean EEG slowing index, calculated by: (−z_F2 − z_F3 + z_F4 + z_F6)/4. **(B)** Residual factor expression after subject-level regression of each factor on the slowing index, averaged within phenotype. A marked cell (*) denotes a deviation beyond the 95th percentile of a max-statistic null from 5,000 phenotype-label permutations across all 84 cells. **(C)** Per-subject distribution of the slowing index by phenotype (half-violin kernel density estimate, Tukey box- and-whisker, individual observations, and group mean [diamond]). *AD, Alzheimer disease; bvFTD, behavioral variant frontotemporal dementia; CBS, corticobasal syndrome; CU, cognitively unimpaired; dAD, dysexecutive Alzheimer disease; DLB, dementia with Lewy bodies; LANS, limbic-predominant amnestic neurodegenerative syndrome; lvPPA, logopenic variant primary progressive aphasia; nfvPPA, nonfluent variant primary progressive aphasia; NPH, normal pressure hydrocephalus; PCA, posterior cortical atrophy; PPAOS, primary progressive apraxia of speech; PSP, progressive supranuclear palsy; SD, semantic dementia*.

DLB MPE demonstrated parietooccipital hypometabolism with relative preservation of FDG metabolism in the posterior cingulate gyrus (i.e., “cingulate island sign”; Fig. 4A). This expression correlated positively with the slowing index (r = 0.51; 95% CI: 0.44-0.57; *P* < 0.001; Fig. 4B). No other class-specific MPE was associated with the slowing index after controlling for DLB MPE (all p > 0.2); results are therefore reported for DLB MPE only. As expected, the LCA placed LANS at the limbic end and dysexecutive AD at the neocortical end (Fig. 4C). Across the cohort, a higher LCA was associated with higher factor 4 (r = 0.44; adjusted for combined dysexecutive AD and LANS MPE, β = 0.34, 95% CI 0.25–0.43, Holm-adjusted *P* < 0.001; Fig. 4D). This association persisted after adjustment for age and DLB MPE (β = 0.21, 95% CI 0.12– 0.31, Holm-adjusted *P* < 0.001) and after further adjustment for phenotype (β = 0.23, 95% CI 0.12–0.35, Holm-adjusted *P* < 0.001). The LCA was also associated with the slowing index and each of the non-artifactual factors; however, none of these associations survived adjustment (Supplementary Fig. 3).

**Figure 4:**
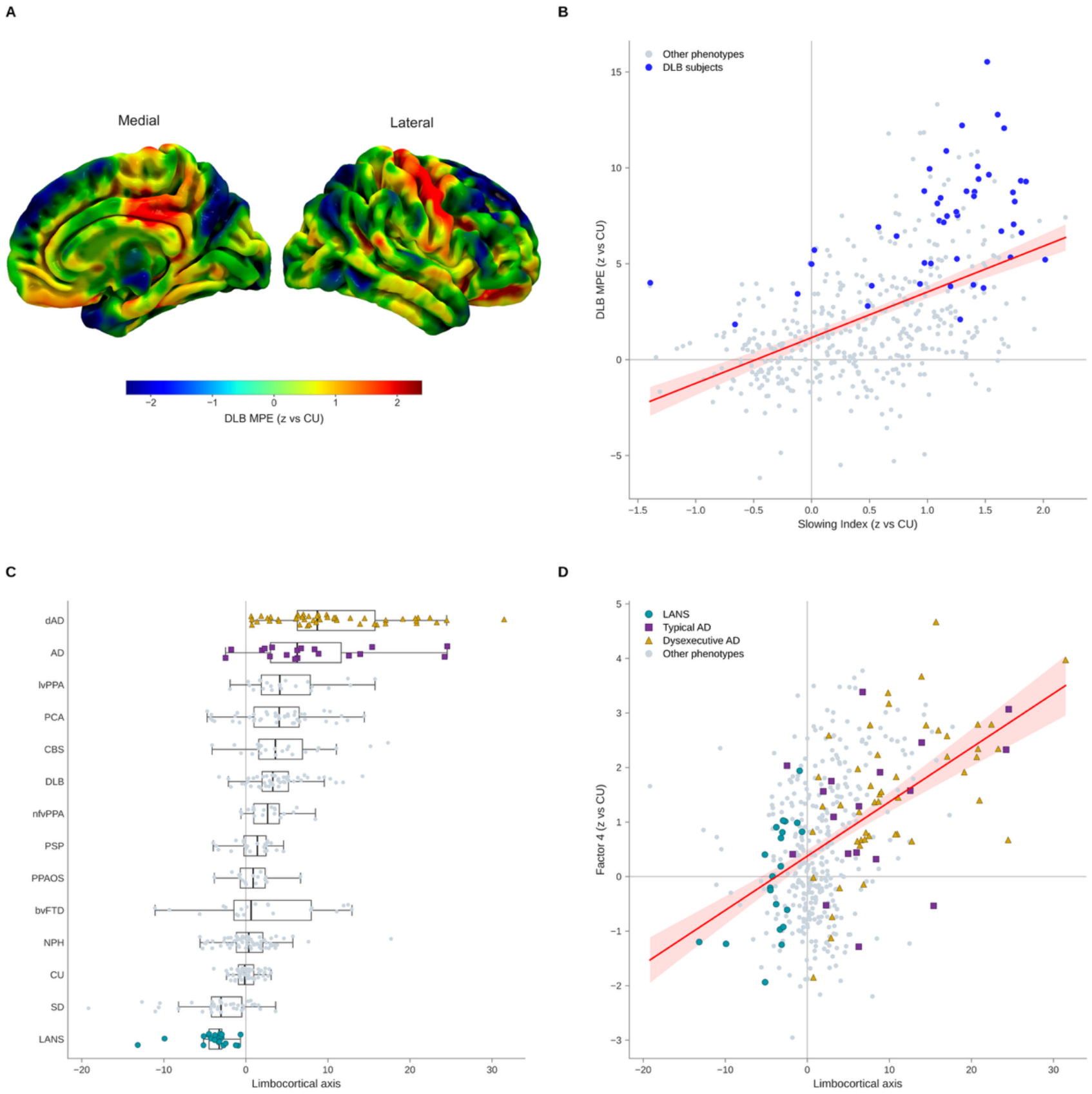
EEG correlation with FDG-PET MPE across phenotypes. **(A)** Group-level FDG-PET MPE for the DLB phenotype rendered on the left hemisphere (medial and lateral views), expressed in standard deviation units relative to the cognitively unimpaired class. Cool colors indicate relative hypometabolism. **(B)** DLB template expression plotted against the EEG slowing index for all subjects (n = 455) with paired EEG factor and FDG-PET template data. Subjects carrying the DLB phenotype label are shown in blue, all other phenotypes in grey. The line is the OLS fit across the full cohort with its 95% confidence band (r = 0.509, *P* < 0.001). **(C)** LCA by phenotype, calculated as dysexecutive AD MPE minus LANS MPE, each standardized against the CU class. **(D)** Factor 4 (frontotemporal delta–theta) plotted against the LCA for all subjects. OLS and 95% confidence band shown in red (r = 0.444, *P* < 0.001). LCA associations with slowing index and factors 2, 3 and 6 are shown in Supplementary Figure 3. *AD, Alzheimer disease; CU, cognitively unimpaired; DLB, dementia with Lewy bodies; LANS, limbic-predominant amnestic neurodegenerative syndrome; LCA, limbocortical axis; MPE, metabolic pattern expression; OLS, ordinary least squares*.

Voxelwise models of FDG uptake on the four neurophysiological factors yielded a distinct metabolic topography for each factor (Fig. 5). Factor 2 was associated with increased metabolic activity in posterior medial and lateral parietal and occipital cortex; its map resembled meta-analytic maps for attention, visual, action, task and spatial (r = 0.33–0.48). Factor 3 was associated with higher metabolism in lateral and medial prefrontal and parietal association cortex decoding to attention, task, numerical, and memory topic-terms (r = 0.31–0.40). Factor 4 was associated with lower metabolism in lateral parietal, precuneus and lateral prefrontal cortex and relatively higher sensorimotor metabolism. Its positive pole decoded to motor, pain, fear and stress topic-terms (r = 0.26–0.37) while its negative pole suggested inverse association with calculation, memory, attention, and language (r = −0.25 to −0.37). Factor 6 was associated with lower medial occipital and relatively higher parietal and precuneus metabolism, decoding to calculation, attention, memory, and language topic-terms (r = 0.17–0.31). For factors 2, 3 and 6, the negative pole decoded to affective and reward terms such as stress, fear, mood, reward and pain. Only the factor 4 map aligned with G1, with lower metabolism toward its transmodal pole (ρ = −0.57, spin-test *P* < 0.01, Holm-adjusted *P* = 0.02). No map aligned with G2 after correction; the nominal factor 4 association (ρ = −0.45, spin-test *P* = 0.031, Holm-adjusted *P* = 0.125) reflected a non-monotonic relationship mirroring that between the two gradients themselves, with correspondence strongest at the somatosensory end of G2 (Supplementary Fig. 4).

**Figure 5:**
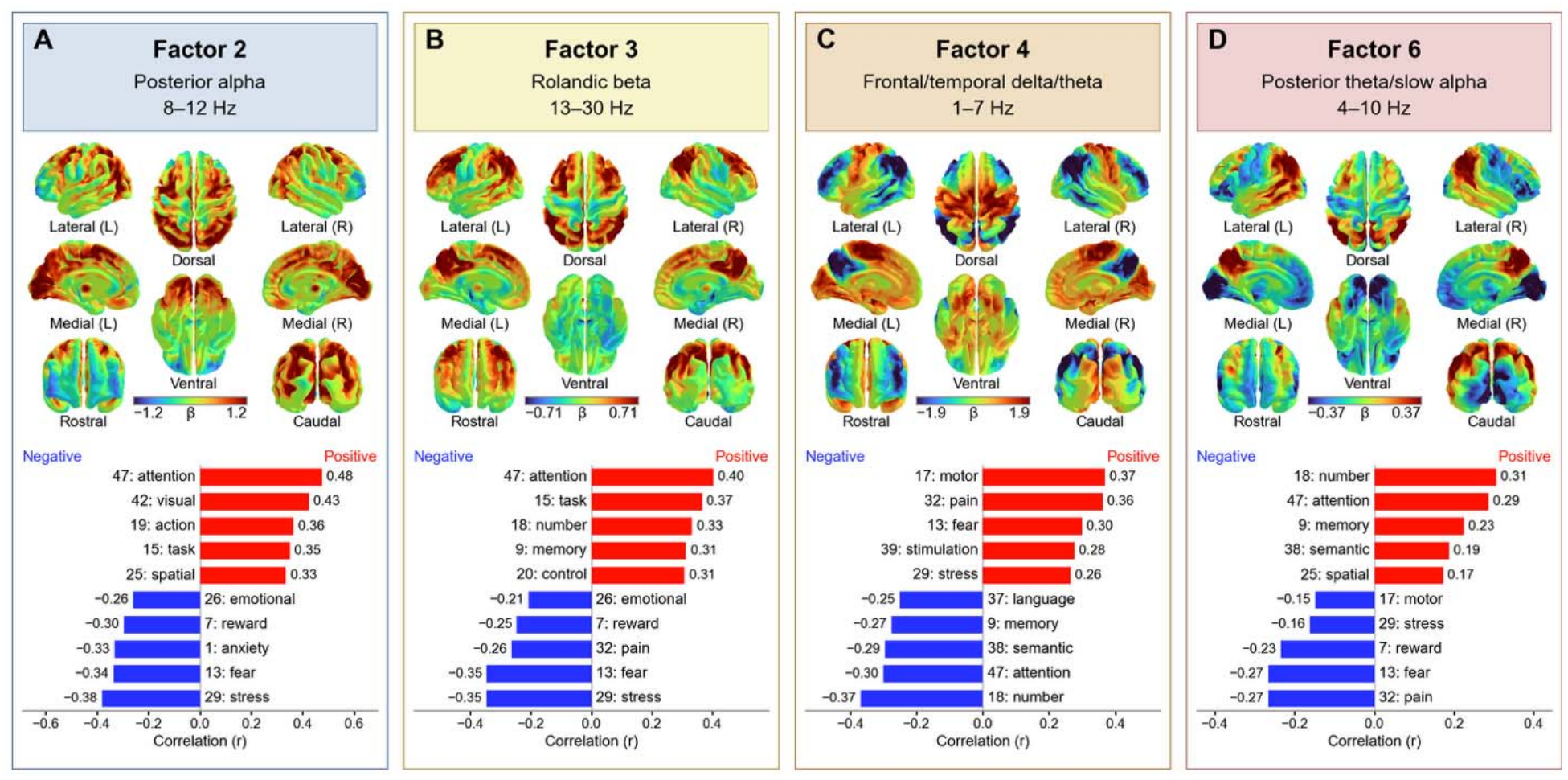
**Functional decoding of the four neurophysiologic EEG factors**. *Upper panels*: Voxelwise regression β maps projected onto the cortical surface in lateral, medial, dorsal, ventral, rostral and caudal views (L, left; R, right). β is the change in FDG SUVR per unit factor score, adjusted for the other three factors. Color scales are symmetric about zero and specific to each factor. *Lower panels*: Neurosynth functional decoding of associated β maps. Top five strongly correlated and strongly anti-correlated topic-terms are presented with their associated family number. *SUVR, standardized uptake value ratio*.

Ninety-eight participants completed neuropsychological testing a median of 23 days (IQR 1–82) from FDG-PET. This subset did not differ from the rest of the cohort in age, sex, years of education, race or phenotype distribution (Supplementary Table 2). After adjustment for age and years of education, higher factor 3 was associated with better attention/executive performance (ρ = 0.33). Increased factor 4 was associated with poorer learning and memory (ρ = −0.33), visuospatial (ρ = −0.38) and global performance (ρ = −0.35; all Holm-adjusted *P* < 0.05; Fig. 6; Supplementary Fig. 5B). Factor 2 showed consistent but non-significant positive associations (ρ = 0.14–0.26), and factor 6 was unrelated to any domain (|ρ| ≤ 0.10). All associations attenuated after adjusting for phenotype, although directions were preserved (factor 3, ρ = 0.09–0.25; factor 4, ρ = −0.18 to −0.28; Supplementary Fig. 5C).

**Figure 6:**
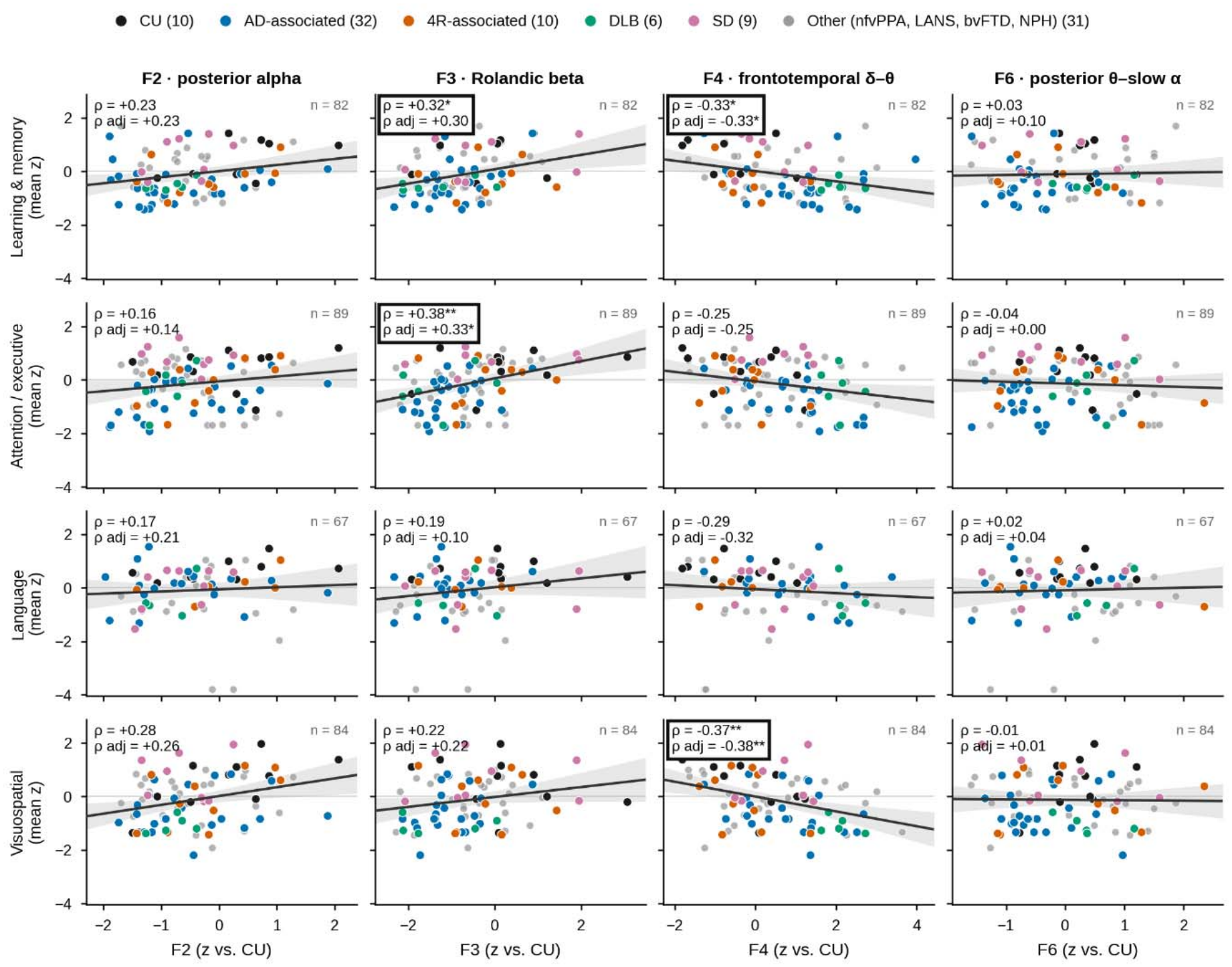
Associations between EEG latent factors and neuropsychological domain performance. Domain composite scores (mean within-test z; higher = better performance) plotted against factor expression (z relative to cognitively unimpaired participants) in the 98 participants with neuropsychological testing. Point color indicates FDG-PET phenotype group (n in parentheses). Lines show ordinary least-squares fits with 95% confidence bands. Each panel reports the unadjusted Spearman ρ, the partial ρ adjusted for age and education (ρ adj), and the number of participants. Bold box and asterisks indicate Holm-adjusted significance across the 20 composite × factor comparisons in each model (\**P* < 0.05, \*\**P* < 0.01). The global composite results appear in Supplementary Figure 5. *LANS, limbic-predominant amnestic neurodegenerative syndrome; bvFTD, behavioral variant frontotemporal dementia; nfvPPA, nonfluent variant primary progressive aphasia; NPH, normal pressure hydrocephalus*.

Using only the four neurophysiological factors, logistic regression discriminated CU from seven of 13 phenotypes after Holm correction (Fig. 7A; Supplementary Table 3). Discrimination was highest for DLB, PCA and dysexecutive AD (AUC 0.85 each; 95% CI 0.76– 0.93, 0.76–0.93 and 0.76–0.92) and typical AD (0.81, 0.67–0.92). AUC was 0.72 (0.65–0.79) for CU versus all impaired participants, 0.82 (0.75–0.89) versus AD-associated and 0.63 (0.53–0.74) versus 4R-associated phenotypes (Holm-adjusted P ≤ 0.011; Fig. 7B). The four-class model reached a macro-averaged AUC of 0.71 (0.67–0.75) and balanced accuracy of 0.47 (chance 0.25; Fig. 7C, D). Lower factor 3 and lower factor 2 contributed most consistently to discrimination from CU (95% CI excluding zero in 8 and 7 of 13 models, respectively). The four-class model relied chiefly on factor 4, higher in AD-associated and DLB and lower in 4R-associated and semantic dementia participants, and on lower factor 6 in AD-associated participants (Fig. 7E). Performance was similar across the five classifiers (mean AUC 0.69–0.72; Supplementary Fig. 6).

**Figure 7:**
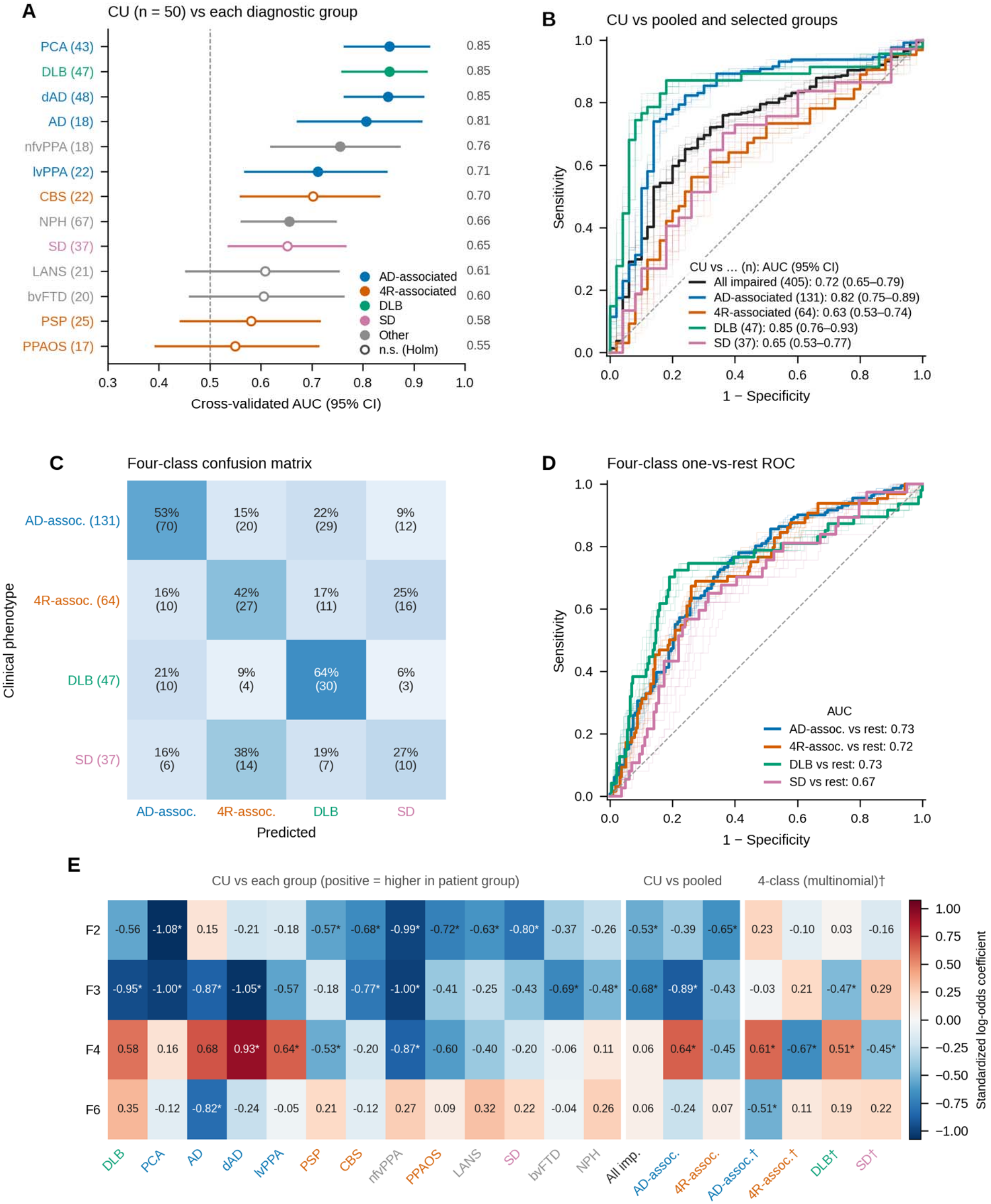
Single-subject classification of clinical phenotypes from four non-artifactual EEG factors. L2-regularized logistic regression was evaluated by ten-times-repeated stratified five-fold cross-validation, with out-of-fold scores averaged across repeats. **(A)** Cognitively unimpaired (CU; *n* = 50) versus each diagnostic group, ordered by cross-validated AUC. Filled markers denote comparisons significant by label-permutation test after Holm correction across the 13 comparisons (*P* < 0.05). **(B)** ROC curves for CU versus all impaired participants (n = 405), the AD-associated group (n = 131), the 4R-associated group (n = 64), DLB (n = 47) and SD (n = 37). Bold curves are derived from repeat-averaged out-of-fold scores, and faint curves show the individual cross-validation repeats. **(C)** Row-normalized confusion matrix for the four-class model separating AD-associated, 4R-associated, DLB and SD participants. Each cell gives the percentage of each clinical phenotype (rows) assigned to each predicted class (columns), with counts in parentheses. **(D)** One-versus-rest ROC curves for each class of the four-class model. **(E)** Standardized logistic regression coefficients from full-data fits for each factor (rows) and group (columns). ^†^Multinomial coefficients represent each group’s deviation relative to the other three groups. *Indicates 95% confidence intervals exclude 0. *AUC, area under the ROC curve; ROC, receiver operating characteristic;* rest abbreviations as in Figure 2.

## Discussion

Across 455 participants spanning 14 FDG-PET-defined phenotypes, four data-driven EEG factors with neurophysiological spectral and spatial signatures varied systematically with metabolic phenotype, whereas the two artifact-dominated factors showed no robust association. Four observations follow. First, much of this variation reflects a single disease-general slowing axis, on which every disease phenotype departed from CU and DLB lay furthest. Second, two factors carried topographic information beyond that axis: frontotemporal delta–theta exceeded what slowing predicted in typical and dysexecutive AD, and posterior theta–slow alpha fell short of it across the AD spectrum, separating DLB from typical AD, dysexecutive AD and PCA. Third, the factors were anchored to biology via regional metabolism and cognitive performance. Fourth, single-subject discrimination was strongest for AD-spectrum and Lewy body phenotypes and weaker for 4R-associated and other frontotemporal phenotypes, marking both the potential and the limits of single-state frequency-based analysis of resting eyes closed EEG as a scalable marker of neurodegenerative network dysfunction across the full dementia spectrum. By anchoring EEG features to FDG-PET–derived phenotypes rather than clinical diagnoses alone, this work clarifies the physiological relevance of specific oscillatory domains and supports a biology-informed framework for EEG-based characterization of neurodegenerative syndromes.

### A Disease-general Slowing Index

Every disease phenotype had a higher mean slowing index than the CU class, from +0.15 standard deviations in progressive supranuclear palsy to +1.10 standard deviations in DLB (Fig. 3A, C). This transdiagnostic shift, an attenuation of faster rhythms with relative augmentation of slower activity, is the most reproducible electrophysiological correlate of neurodegeneration.^14,15,40,41^ A shared axis of this kind is consistent with dimensional models in which neurodegenerative phenotypes occupy positions along common continua of network dysfunction rather than discrete categories.^1,2^ Adjusting for this axis showed how much of each factor’s phenotypic structure it explains. Phenotype-attributable variance fell by more than half for posterior alpha (η² 0.125 to 0.047) and Rolandic beta (0.154 to 0.035) and by nearly half for frontotemporal delta–theta (0.216 to 0.118), whereas it rose for posterior theta–slow alpha (0.082 to 0.147; Fig. 3B). Between-phenotype differences in alpha and beta therefore largely reflect how much a syndrome is slowed overall, whereas the slow-wave factors also carry information about where the slowing is expressed. Given that posterior alpha and posterior theta–slow alpha share a posterior topography and occupy adjacent frequency ranges, this finding may represent that slowing of the posterior dominant rhythm shifts spectral power from the former toward the latter.^42,43^ Because each factor is also one component of the slowing index, part of its variance is removed by construction; the residuals are therefore conservative and not fully independent of one another.

It should be noted that the DLB phenotype, which showed the most overall slowing, had the largest absolute value deviations from CU, yet minimal residuals after removing slowing (all |residual| ≤ 0.15, Fig. 3B). Amongst all the neurodegenerative syndromes, DLB diagnostic criteria are the only major neurodegenerative diagnostic criteria that include EEG as a supportive biomarker.^9^ However, the findings here suggest that the previously reported quantitative EEG approaches are oversimplifications of the complexity of multi-regional dysfunction that is embedded within the disease.^43–46^ The DLB metabolic pattern expression reproduces parietooccipital hypometabolism with relative preservation of the posterior cingulate (i.e., the cingulate island sign), a pattern with recognized diagnostic specificity rather than a generic index of disease burden (Fig. 4A). The EEG slowing index correlated with DLB template expression across all subjects irrespective of assigned phenotype label (Fig. 4B). Hence, oscillatory features isolated by tensor decomposition track disease biology rather than clinical severity alone: the same EEG measure that separates phenotypes also varies continuously with a biologically anchored marker of the underlying disease process.

### Oscillatory Interpretation Beyond the Slowing Index

Factor 2, corresponding to posterior alpha activity, demonstrated consistent reductions in DLB and PCA (Fig. 2A and 3A). The mechanisms underlying this reduction are likely different for each of these pathologies. Posterior alpha rhythms arise from coordinated thalamocortical and corticocortical interactions and reflect the functional integrity of large-scale posterior association networks rather than a passive “idling” state.^47,48^ Rhythms generated in the occipitoparietal cortex are propagated to the pulvinar where alpha synchrony regulates between visual areas according to attention demands.^47,48^ Although both syndromes present with prominent visuospatial and visuoperceptual deficits,^9,24^ the residual analysis suggests that alpha diminishment may occur variably. In DLB, alpha loss was fully accounted for by overall slowing (residual +0.08; Fig. 3B). Such a pattern, where posterior alpha slows into the alpha-theta range, is a known hallmark of the condition.^9,44^ Alpha reactivity and its attentional modulation are shaped by basal forebrain cholinergic input.^49,50^ In both DLB and AD, reduced alpha reactivity tracks atrophy of the nucleus basalis of Meynert and degeneration of its cholinergic pathways.^51,52^ Interestingly, in both AD and dysexecutive AD, alpha loss was also accounted for by overall slowing (Fig. 3A and 3B). PCA, in contrast, had posterior alpha loss that exceeded that predicted by slowing (residual −0.31) and this was not offset by posterior theta-slow alpha (residual −0.45, Fig. 3B). Notably, posterior alpha carried the largest coefficient of any factor in separating PCA from the CU class (Fig. 7E). Such a pattern is more consistent with the loss of alpha generators themselves rather than slowing of an intact rhythm. In this AD phenotype, posterior tau deposition is thought to be the primary driver of symptoms, with nearly all patients with this diagnosis having a tau-positive PET scan.^53^ The early deposition of tau in the primary visual cortex and the occipitoparietal junction could thus cause breakdown of alpha rhythm, which is believed to have dominantly supragranular generators in this area of the cortex^47,54^ and infragranular generators in others.^55,56^ Factor 2 may therefore index the integrity of posterior association networks, with its loss in PCA plausibly reflecting local degeneration of alpha generators and its loss in DLB, and other AD phenotypes, possibly reflecting cholinergic dysfunction, including disrupted septo-hippocampal regulation of cortical oscillatory dynamics.^13^

Factor 3, corresponding to Rolandic beta activity, was notably reduced, compared to the CU group in dysexecutive AD, PCA, and DLB. It also separated the latter from several other non-amyloid related phenotypes (Fig. 2B). Beta oscillations are a core mechanism supporting long-range cortico-cortical communication within frontoparietal control networks and are critically involved in the top-down executive control, integration of sensorimotor processes, visuo-perceptual processing, and working memory. Rather than merely preserving the current cognitive state, beta synchronization flexibly engages transient network interactions within and between cortical areas, enabling flexible ensemble formation needed to uphold internal task demands when external stimuli are absent or when task rules must be internally maintained.^57,58^ Rolandic beta was the most closely tied to the slowing index as adjustment removed more than three-quarters of its phenotype-attributable variance and left no phenotype residual larger than 0.23 (Fig. 3B). In addition, reduced Rolandic beta reliably distinguished the most phenotypes (eight out of 13) from CU (Fig. 7E). The reduction of Rolandic beta in dysexecutive AD adds to previously reported selective hypometabolism on FDG-PET of parieto-frontal brain regions in providing evidence that dysexecutive AD represents a biologically distinct subtype of AD.^21^ In DLB, the reduction in Rolandic beta could be biologically correlated with this oscillation’s known strong cholinergic association and the disease’s reduced cholinergic projections.^50^ In Parkinson disease, there is a hypercholinergic state in the striatum leading to a lack of attenuation of Rolandic beta during contralateral motor activation, suggesting a possible role of EEG in distinguishing between the two conditions which often have phenotypic overlap.^59^ Finally, the relative reduction of this factor in PCA can be best explained by the known role of beta oscillations in the posterior parietal lobe in visual perception and processing.^58^

Factor 4, representing frontotemporal delta-theta (slow-wave) activity, was markedly elevated in dysexecutive AD and DLB relative to multiple other phenotypes (Fig. 2C). Slow wave activity in the frontal cortex has multiple generators including thalamic nuclei and layer V cortical neurons; however, in the context of resting-state EEG in neurodegeneration, should be interpreted as the loss, or disruption, of other higher frequency (e.g., alpha, beta) bands that are more typically seen.^60^ Hence, it is not surprising that syndromes that are elevated in factor 4 are also the ones with reduced factor 3 (Rolandic beta, Fig. 3B). In dysexecutive AD, the prominence of frontally distributed slow-wave activity is biologically consistent with selective vulnerability of frontoparietal executive networks, which are disproportionately affected in this phenotype compared to typical amnestic AD.^21^ While this is supported by this phenotype having the strongest coefficient in our predictive algorithm, factor 4 was also elevated in all AD-associated phenotypes (Fig. 3B, Fig. 7E). These findings may hence support a previous report that delta–theta hypersynchrony in frontotemporal lobes colocalizes with both amyloid-beta and tau deposition, known biomarkers of AD.^61^ In DLB, increased slow-wave activity may arise from combined cholinergic degeneration and thalamocortical dysrhythmia, both of which are known to shift oscillatory balance toward theta dominance.^51^ Quantitative EEG findings suggest that frontotemporal slowing may be more pronounced in DLB than AD;^62,63^ however, the dominance of factor 4 in AD-phenotypes is suggestive that this latent factor encodes syndrome-specific network failure involving executive and attentional circuitry and is not purely dependent on numerical quantification of frequency.

Factor 6, corresponding to posterior theta and slow alpha activity, is increased in both DLB and normal pressure hydrocephalus compared to both typical AD and dysexecutive AD (Fig. 2D). Theta slowing is postulated to be due to thalamocortical dysrhythmia, and when found in the posterior region, it is a well described biomarker that distinguishes DLB from typical AD.^46,51,64^ Conversely, factor 6 also captured the AD spectrum, through its reduction. It was the only factor whose phenotype-attributable variance increased after adjustment for overall slowing (η² 0.082 to 0.147), and typical AD produced the only residual to exceed the family-wise threshold, with less posterior theta–slow alpha than its slowing predicted (Fig. 3A and 3B).

Dysexecutive AD and PCA deviated in the same direction, and lower factor 6 characterized AD-associated participants in the four-class model (Fig. 7E). Read together with factor 2, this distinguishes slowing of the posterior rhythm from its loss. In DLB, posterior alpha declines as theta–slow alpha rises, consistent with a slowed but intact generator. In typical AD, dysexecutive AD and PCA, both decline, consistent with tau-related degeneration of the posterior association cortex that generates the rhythm.^53^ A reduction in factor 6 may therefore mark AD-spectrum posterior network failure rather than simply the absence of Lewy body physiology. The increase in factor 6 in normal pressure hydrocephalus is also biologically plausible, as ventricular enlargement and periventricular white matter disruption are known to impair long-range connectivity and promote diffuse slowing on EEG.^39,65^ Increase in posterior theta power, in particular, has been correlated with worse gait dysfunction in this disease.^65^ Finally, it should be noted that semantic dementia had an increase in factor 6 compared to typical AD. These findings, together with the classification results, contrast with the traditional framework that semantic dementia does not have changes in routine EEG.^66–68^

### Neurobiological anchoring of EEG factors

The EEG factors were further anchored to neurobiology by linking each to the regions whose metabolism it tracks, ^69^ to the cognitive functions those regions support, inferred by meta-analytic decoding across the Neurosynth database of functional neuroimaging studies, and to measured cognitive performance. Voxelwise regression, with the four factors modeled jointly, mapped each factor onto a distinct metabolic topography (Fig. 5). For posterior alpha and Rolandic beta, which reflect physiological rhythms, meta-analytic decoding matched each rhythm’s known function; for the two slow-wave factors, which index dysfunction, the maps instead resembled known disease patterns. Posterior alpha was associated with posterior parietal and occipital metabolism and decoded to visual, attentional and spatial processing, consistent with the role of alpha in gating visual attention.^48,70^ Rolandic beta, despite its central scalp topography, mapped not onto the sensorimotor strip but onto lateral and medial prefrontal and parietal association cortex, decoding to attention, task and control-related terms, in keeping with EEG–fMRI evidence that resting beta power covaries with default-mode and frontoparietal network activity.^71,72^ The frontotemporal delta–theta map resembled the heteromodal hypometabolism with relative sparing of primary cortices that characterizes AD.^73^ The map’s sparing of the mesial temporal lobe may explain the low expression of this factor in LANS cases (Fig. 2C and 7E). Supporting this interpretation, factor 4 tracked the LCA at the level of individual patients (Fig. 4D). Because the index derives from phenotype templates rather than the voxelwise maps, this supports the view that factor 4 reflects neocortical rather than limbic dysfunction. This pattern also motivated quantitative comparison to the principal gradient of cortical organization (G1),^32^ which our group previously demonstrated correlation with the second eigenbrain of the global functional state space.^1^ Of the four factor maps, only factor 4 was significantly aligned with G1 (Supplementary Fig. 4). Hence, this factor map indexes how far a patient’s metabolism has shifted along this macroscale hierarchy that fails early in AD^73^ rather than a focal frontotemporal abnormality alone. Finally, the posterior theta–slow alpha map reproduced the medial occipital hypometabolism with relative posterior cingulate sparing that distinguishes DLB from AD.^74–76^ Apart from the gradient comparison, these analyses are descriptive rather than inferential. Neurosynth decoding relies on reverse inference from task activation in healthy participants and therefore suggests rather than establishes the cognitive functions involved,^29,30^ and because FDG uptake is expressed relative to a reference region, regions spared by disease can appear relatively hypermetabolic.^77^

The 98 participants with neuropsychological data were reasonably representative of the entire cohort (Supplementary Table 2). Adjusting for age and education, factor 3 correlated with better attention/executive performance (Fig. 6 and Supplementary Fig. 5B). This accords with the proposed role of beta in maintaining and reactivating task-relevant content through long-range frontoparietal coordination.^57,58^ Factor 4 correlated with poorer learning and memory, visuospatial, and global performance (Fig. 6 and Supplementary Fig. 5B). Its metabolic correlate explains this profile. Higher frontotemporal delta–theta was associated with lower metabolism in lateral parietal cortex and precuneus, regions central to episodic memory retrieval and visuospatial processing,^78,79^ and in lateral prefrontal cortex, which together make up much of the heteromodal association cortex that fails early in AD.^73^ After further adjustment for phenotype, reported associations attenuated, although each kept its direction (Supplementary Fig. 5C), suggesting that these neuropsychological associations are driven largely by between-phenotype differences. Posterior theta–slow alpha shows why this matters. As noted above, factor 6 is increased in DLB yet reduced in AD-phenotypes, yet both groups are cognitively impaired, so when all participants are pooled these opposite shifts cancel and the factor appears unrelated to cognition. Within-phenotype estimates were unfortunately underpowered with this dataset.

### An oscillatory account of the phenotypes

The SLOD framework treats the hippocampus as a generative engine that conditions neocortical activity, with the coupling between the two scheduled by rhythmic activity that scalp EEG samples directly.^13^ On this account, the dementias are not a set of independent regional lesions but failures at different points along a hippocampal–neocortical axis, and phenotypes that differ in where along that axis they fail should differ in their resting oscillatory dynamics. We tested this directly with the LCA. Derived from the FDG-PET templates rather than from the EEG, it placed LANS at the limbic pole and dysexecutive AD at the neocortical pole, and frontotemporal delta–theta rose with it across all 455 participants, within phenotype as well as between phenotypes, and after adjustment for age and for DLB expression (Fig. 4C, D). Factor 4 was also the only map aligned with the principal gradient of cortical organization. Together, these findings support the interpretation of slow frontotemporal activity as an electrophysiological readout of degeneration along the limbic–neocortical axis, with its associated metabolic topography organized along the principal gradient from unimodal to transmodal cortex. This convergence links Factor 4 to the hierarchical functional physiology that the SLOD framework predicts should shape the oscillatory regime.

DLB offers a complementary test. Its slowing was the most severe of any phenotype yet carried no topographic residual, and the slowing index tracked DLB MPE continuously across the cohort. In SLOD terms, this is the signature of a disrupted scheduler rather than a damaged decoder: cholinergic septo-hippocampal and thalamocortical inputs that set the pace of hippocampal–neocortical coupling fail, slowing the posterior rhythm into the theta range while leaving its generators, and the neocortical targets they condition, relatively intact, which is also why cingulate metabolism is relatively preserved while posterior cortex is hypometabolic (Fig. 4A). The framework thus assigns distinct computational failure modes to the two phenotype families this study resolved best and predicts that the frontotemporal dementia-spectrum syndromes it resolved poorly (e.g, progressive supranuclear palsy and primary progressive apraxia of speech), whose pathology falls outside the hippocampal–neocortical axis, would require measures of dynamics rather than resting spectral power.

### Predictive capacity of EEG factors

Using only the four neurophysiological factors, logistic regression was able to distinguish AD-phenotypes, nonfluent primary progressive aphasia, DLB, and normal pressure hydrocephalus from the CU class (Fig. 7A). In four-class analysis, DLB was the best recalled class and one-vs-rest AUCs were between 0.67 and 0.73 (Fig. 7C and 7D). Overall performance was similar to previous efforts,^17,46,80^ but this approach advances on prior efforts in several aspects of study desig. Phenotypes were defined by FDG-PET metabolic patterns rather than by clinical diagnosis alone; the features were four physiologically interpretable factors, derived without reference to diagnosis and separated from artifact, rather than large sets of band-power or connectivity measures selected for their discriminative value; and the analysis spanned 13 phenotypes, including atypical AD variants and rarer frontotemporal dementia syndromes, rather than the two-group comparisons typical of prior work. Mean AUC was similar across five algorithms of very different flexibility (Supplementary Fig. 6), indicating that the discriminative information lies in the factors themselves rather than in the choice of model.

### Limitations

Several limitations warrant consideration. FDG-PET derived phenotypes were used as the reference standard rather than neuropathologic confirmation, although prior work has demonstrated strong correspondence between StateViewer classifications and underlying pathology.^5^ Medications (e.g., cholinesterase inhibitors, benzodiazepines) and comorbidities (e.g., epilepsy) known to alter the resting EEG were not accounted for and may have influenced factor expression. The cohort was predominantly non-Hispanic White, which may limit generalizability to more diverse populations. There is also selection bias inherent in a cohort of convenience. While the interpretable nature of the analysis supports biologically plausible inferences linking clinical syndromes to oscillatory patterns, these relationships were not formally tested here. Subsets of this cohort have additional cognitive and imaging data available, and direct association of latent factor levels with these measures is a focus of ongoing work.

Finally, this study focused on resting-state EEG during eyes-closed wakefulness. Reactivity, temporal variability, rhythmicity,^81^ aperiodic activity,^82^ as well as task-based and longitudinal recordings, may reveal additional disease-relevant dynamics especially relevant to the physiology of 4R-tau associated degenerative syndromes.

## Conclusion

Data-driven decomposition of routine clinical EEG isolates oscillatory components that are separable from artifact and that track FDG-PET–defined phenotype, regional metabolism and cognition. Much of this information lies on a shared slowing axis, carried chiefly by the loss of Rolandic beta and posterior alpha, on which every disease phenotype had a higher mean value than CU. Beyond that axis, the slow-wave factors distinguish how syndromes fail: frontotemporal delta–theta tended to exceed what slowing predicted in typical AD and dysexecutive AD, posterior theta–slow alpha fell short of it in typical AD, dysexecutive AD and PCA and separated DLB from each, and posterior alpha loss may reflect local degeneration in PCA but slowing of the rhythm in DLB. These findings fit models in which cognition depends on the oscillatory coordination of large-scale brain systems, including hippocampal–neocortical interactions, and in which the major dementias reflect selective failures of that coordination.^1,2,13^ While this work is most informative about AD-phenotypes and DLB, future work is needed to uncover neurophysiological associations in frontotemporal dementia-spectrum phenotypes and to correlate findings with other neuroimaging, cerebrospinal fluid-based, and blood-based biomarkers.

## Acknowledgements

The authors thank the participants and their families for their contributions to the Mayo Clinic Alzheimer’s Disease Research Center, the Mayo Clinic Study of Aging and the Neurodegenerative Research Group, and the patients of the Mayo Clinic neurology practice who authorized use of their health records for research. We thank the staff of the Mayo Clinic Department of Neurology EEG Laboratory and the Division of Nuclear Medicine for acquisition of the clinical EEG and FDG-PET studies. During manuscript preparation, the authors used large language models to assist with language editing and improving clarity and readability. The authors reviewed and edited all AI-assisted text and take full responsibility for the content of the manuscript.

## Funding

This work was funded in part by National Institutes of Health (NIH) grants (P30 AG062677, U01 AG006786, and U01 AG082350-01A1). N.C.L.’s work on this effort is supported by the NIH (K99AG097974-01A1) and the Brain and Behavior Research Foundation (Young Investigator Grant #34277). G.S.D.’s work is supported by NIH (R01AG089380, U01AG057195, U01NS120901, U19AG032438, P30AG062677).

## Competing Interests

D.T.J. is co-founder of Cephlodyne Neurotechnologies and is an inventor on patents owned by Mayo Clinic related to the StateViewer technology used in this study. B.H.B. has licensed IP to Cadence Neuroscience and Seer Medical Pty and has received honoraria and travel support from Eisai Inc. and UCB. He has received research support from Neurelis Inc., UNEEG Medical A/S, and Seer Medical. V.J.L. reports grant support to his institution from Siemens Medical Imaging and Avid Radiopharmaceuticals and honoraria from PeerView Institute for Medical Education. V.J.L. and G.S.D report research materials or services provided to their institution by Avid Radiopharmaceuticals. G.S.D is a co-Project PI for a clinical trial in anti-NMDAR encephalitis, which receives support from Amgen Pharmaceuticals. All other authors have no competing interests.

## Data Availability

The data that support the findings of this study are available from the corresponding author, upon reasonable request.

