## Supplementary Material for "A shared slowing axis and phenotype-specific rhythms link clinical EEG to FDG-PET dementia phenotypes"

**Supplementary Methods**

**Spatial Correspondence to Macroscale Functional Gradients**

To test whether the metabolic topography associated with each EEG factor followed the macroscale organization of the cortex, each of the four β maps from the voxelwise multivariable model was compared with the first two functional connectivity gradients.^1^ The gradients were derived by diffusion map embedding of group-averaged resting-state functional connectivity from the Human Connectome Project. The principal gradient (G1) spans from primary sensorimotor to transmodal default-mode cortex, and the second gradient (G2) separates somatomotor from visual cortex. Both were obtained in fsLR 32k surface space.^2^

Each β map was projected from MNI onto the fsLR 32k surface by registration fusion, with linear interpolation.^3^ Factor maps and gradients were then parcellated with the Schaefer 400-parcel, 7-network atlas by averaging vertex values within each parcel.^4^ Parcellation was used to reduce vertex-level sampling noise from the volume-to-surface projection and to make the spin test computationally tractable. Because the atlas covers cerebral cortex only, subcortical, cerebellar and medial-wall data did not contribute. Because the sign of a diffusion map embedding is arbitrary, the polarity of each gradient was confirmed from its mean value across the most posterior 5% of cortical vertices on the fsLR midthickness surface, corresponding to the occipital pole. This value was negative for G1 (−3.29) and positive for G2 (+3.97). G1 therefore increases from unimodal toward transmodal cortex, and G2 from somatomotor toward visual cortex.

Spatial correspondence between each factor map and each gradient was quantified as Spearman's rank correlation across parcels. Because neighboring parcels are not statistically independent, significance was assessed against a null model that preserves spatial autocorrelation (spin test).^5^ Parcel centroids on the fsLR spherical surface underwent 10,000 random rotations, applied to one hemisphere and mirrored to the other. After each rotation, every parcel took the value of the parcel whose rotated centroid fell nearest to it. The same rotations with a fixed random seed were applied to all four factor maps. Observed and rotated maps were rank-transformed identically before correlation. Two-sided P values were computed as (1 + the number of null correlations whose absolute value equaled or exceeded the observed absolute value)/(10,000 + 1). P values were Holm-corrected across the four factors within each gradient.

**Machine Learning Analysis**

**Classifiers**

We compared five classifiers representing generative, linear, kernel-based and tree-ensemble approaches. The first was Gaussian naïve Bayes, as used in our prior work. The second was L2-regularized logistic regression with an inverse regularization strength of C = 1, fitted with the lbfgs solver. The third was a support vector machine with a radial basis function kernel (C = 1; γ = 'scale'). The fourth was a random forest of 300 trees, with a minimum of 3 samples per leaf and √p candidate features considered at each split. The fifth was histogram-based gradient boosting, with a maximum tree depth of 3, a learning rate of 0.05, 150 boosting iterations and a minimum of 10 samples per leaf. Hyperparameters were fixed *a priori* at conventional values and were not tuned. This choice avoids the optimistic bias that data-driven model selection introduces in samples of this size, and it means that any differences between models reflect the algorithms rather than differential tuning.

**Class-imbalance**

Group sizes were unequal, most markedly in the CU versus all-impaired analysis (50 vs 405). Every classifier was therefore configured so that each class contributed equally to model fitting. Logistic regression, the support vector machine and gradient boosting used balanced class weights, inversely proportional to class frequency. The random forest used balanced class weights recomputed within each bootstrap sample, and naïve Bayes used uniform class priors. As a result, the default decision rule corresponds to the class-balanced threshold: a predicted probability of 0.5 for binary models, and the maximum predicted class score for the four-class model. For the support vector machine, scores were derived directly from the decision function rather than from Platt-scaled probabilities.^6^ Platt scaling is fitted on unweighted data and would shift the decision boundary back toward the majority class. Binary decision values were transformed with a logistic function, and four-class one-versus-rest decision values with a softmax function. Both transformations are monotonic, so they leave the area under the receiver operator characteristic curve (AUC) unchanged and map the decision boundary to the class-balanced threshold.

**Cross-validation**

Performance was estimated by stratified five-fold cross-validation, which preserves class proportions in every fold. The procedure was repeated 10 times with different random fold assignments generated from fixed random seeds. Feature standardization (z-scoring) was estimated on the training folds only and then applied to the held-out fold, so no information from test participants entered model fitting. Each participant therefore received one out-of-fold predicted score per repeat. These scores were averaged across the 10 repeats to give a single cross-validated score per participant, and all performance metrics were computed from these averaged scores. The variability of the AUC across repeats was also recorded and is reported in Supplementary Table 3.

**Performance Metrics**

For the binary analyses, the primary metric was the AUC. We also computed sensitivity, specificity and balanced accuracy (the mean of sensitivity and specificity) at the class-balanced threshold. For the four-class analysis, we report the macro-averaged one-versus-rest AUC, defined as the unweighted mean of the four class-versus-rest AUCs. We also report balanced accuracy, defined as the mean per-class recall, for which chance is 0.25. Finally, we report the row-normalized confusion matrix, in which each participant was assigned to the class with the highest cross-validated score. Ninety-five percent confidence intervals for the AUC were obtained by stratified bootstrap resampling of participants: 2000 resamples were drawn with replacement within each class, and the interval was taken from the 2.5th and 97.5th percentiles of the resulting distribution.

**Model Selection**

L2-regularized logistic regression was designated the primary model for three reasons: it is linear, it yields directly interpretable coefficients. In addition, no alternative classifier materially outperformed it; mean AUC across all analyses ranged from 0.69 to 0.72 across the five algorithms.

**Significance**

Statistical significance for the primary model was assessed by label-permutation testing with 1000 permutations per analysis. Under each permutation, the class labels were randomly shuffled and the complete cross-validation procedure was re-run, including fold assignment, standardization and model fitting. The test statistic was the mean AUC (macro-averaged one-versus-rest AUC for the four-class model) across two repeats of stratified five-fold cross-validation; this reduced number of repeats was used for computational tractability, so the test statistic can differ slightly from the reported AUC. The same statistic was computed on the observed labels using the same fold seeds. The permutation P-value was calculated as one plus the number of permuted statistics greater than or equal to the observed statistic, divided by one plus the number of permutations. Because the entire pipeline is re-run under each permutation, this test accounts for the dependence between cross-validation folds, which violates the independence assumption underlying standard parametric tests of cross-validated AUC. P-values were corrected for multiple comparisons with the Holm–Bonferroni procedure, applied separately within analysis (1) (13 tests) and across analyses (2) to (5) (4 tests). With 1000 permutations, the smallest attainable unadjusted P-value is approximately 0.001.

**Factor contributions**

To characterize the direction and magnitude of each factor's contribution, we fitted L2-regularized logistic regression models (C = 1, balanced class weights) to the full data for each analysis. All four factors were standardized to zero mean and unit variance. For binary models, each coefficient represents the change in the log-odds of membership in the patient group, relative to CU, per standard deviation increase in that factor, conditional on the other three factors. For the four-class model, the multinomial coefficients represent each group's deviation in log-odds relative to the other groups. Uncertainty in the coefficients was estimated by stratified bootstrap with 1000 resamples, and coefficients whose 95% percentile interval excluded zero were flagged. Coefficients are expressed on the raw factor loadings. Because the factors are partially correlated, each coefficient reflects that factor's unique contribution given the others and should be interpreted as descriptive rather than causal.

**Supplementary Figures**

**Supplementary Figure 1: Pairwise comparisons of latent EEG factor scores for Factors 1 and 5 across clinical syndromes.** Each panel shows all pairwise comparisons of subject-level scores for one latent EEG factor among the 14 clinical groups: **(A)** Factor 1 and **(B)** Factor 5. Comparisons are presented as 14×14 matrices, with the upper triangle displaying signed t statistics (row minus column comparisons). No pairwise comparison for either factor remained significant after Holm correction. The color scale is identical to that of Figure 2, so t statistics are directly comparable across all six factors. Abbreviations as in Figure 2.

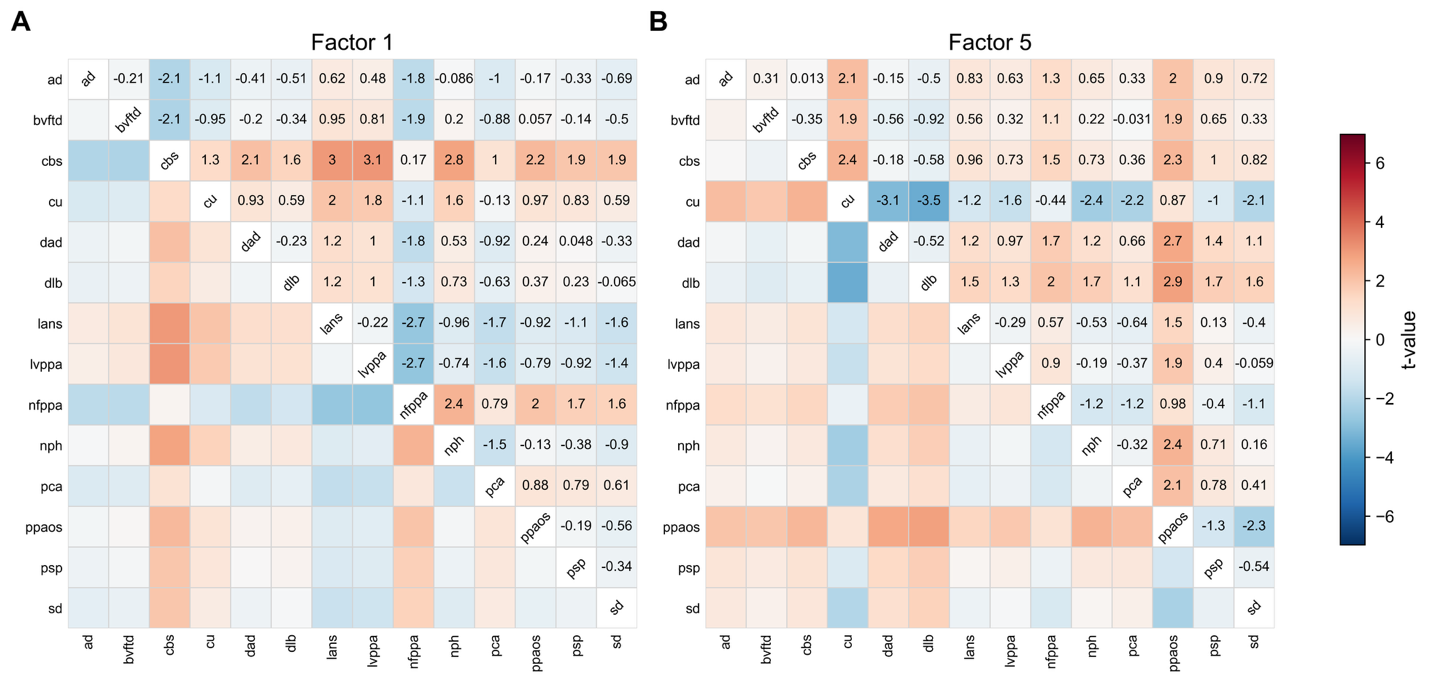

**Supplementary Figure 2: Pairwise group differences in EEG latent factors are stable across the FDG-PET class-confidence threshold.** Each panel compares the primary cohort (FDG-PET class assignment at *p* < 0.01; *n* = 455) with the full temporally matched cohort (*p* < 0.5; *n* = 875) for one EEG latent factor: (**A**) Factor 2, (**B**) Factor 3, (**C**) Factor 4, (**D**) Factor 6. **Upper triangle:** change in the magnitude of the pairwise *t* statistic between cohorts, Δ|*t*| = |*t*|_p<0.01_ − |*t*|_p<0.5_, printed in each cell and shaded on a diverging scale centered at zero. **Lower triangle:** Holm-adjusted significance in each cohort, shown as *p* < 0.01 cohort | *p* < 0.5 cohort, with ns denoting a non-significant comparison. Cell shading indicates concordance: black with white text, significant in both cohorts; dark grey with white text, significant only at *p* < 0.01; light grey with black text, significant only at *p* < 0.5; white, significant in neither. Of the 45 comparisons significant in the primary cohort, 37 are also significant in the full cohort, 8 are significant only in the primary cohort, and 23 are significant only in the full cohort; no contrast reverses direction in either cohort. Abbreviations as in Figure 2.

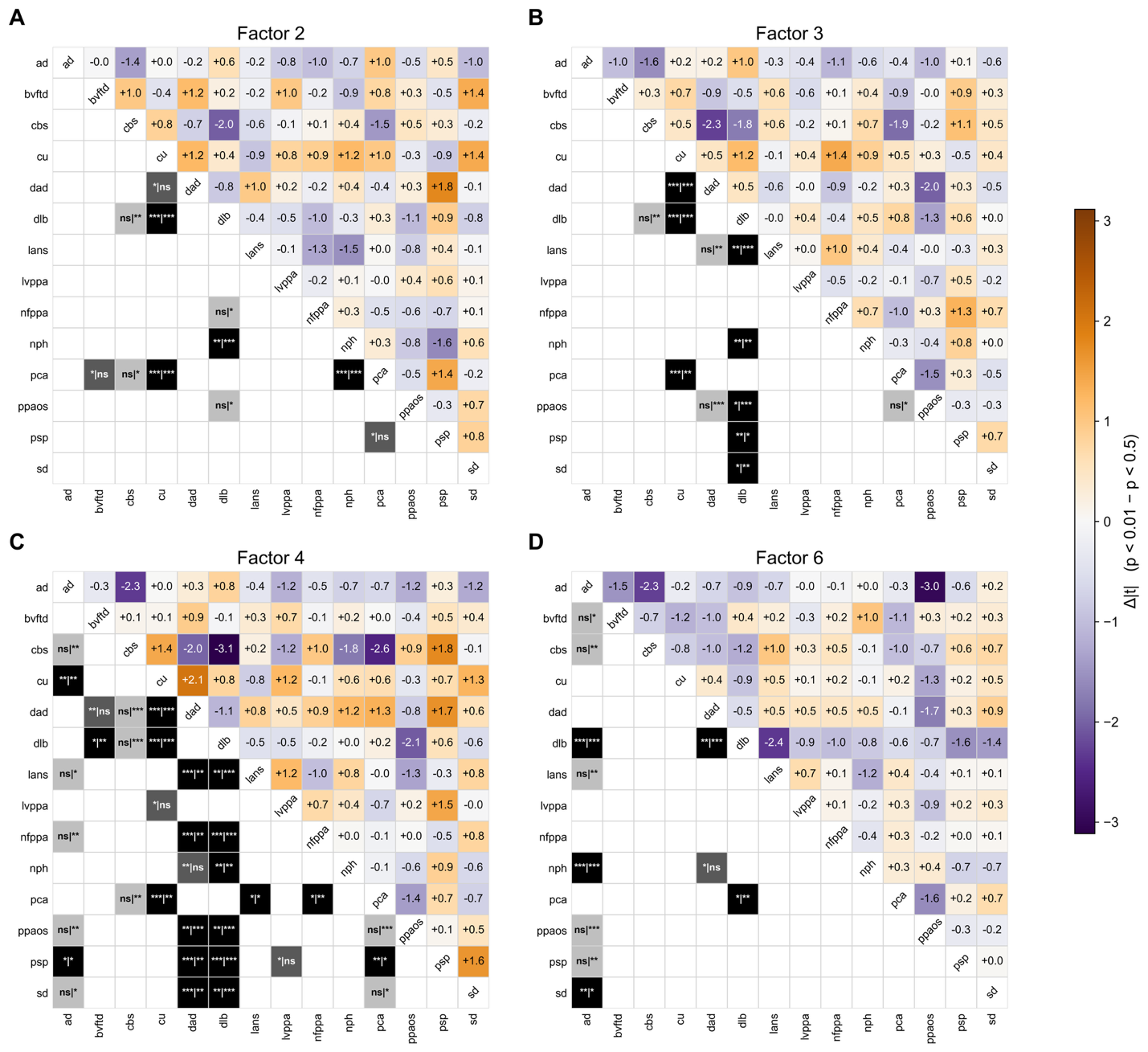

**Supplementary Figure 3: Associations of the neocortical–limbic index with the slowing index and the remaining non-artifactual factors.** The **(A)** slowing index, **(B)** factor 2 (posterior alpha), **(C)** factor 3 (Rolandic beta) and **(D)** factor 6 (posterior theta–slow alpha) plotted against the LCA for all subjects (n = 455). LANS, typical AD and dysexecutive AD subjects are highlighted as indicated in the key, and all other phenotypes are shown in grey. Lines are unadjusted OLS fits with 95% confidence bands. Inset values give the Pearson r with its 95% confidence interval and the standardized LCA coefficient (β) from three nested models: Model 1, adjusted for combined dysexecutive AD and LANS MPE; Model 2, additionally adjusted for age and DLB MPE; Model 3, additionally adjusted for phenotype (14-level fixed effect). P values are Holm-adjusted across the five outcomes (slowing index and factors 2, 3, 4 and 6) within each model; factor 4 is shown in Figure 4D. Factors are standardized against the CU class and shown in their native sign; lower values of factors 2 and 3 and higher values of factor 6 indicate the pathological direction. Abbreviations as in Figure 4.

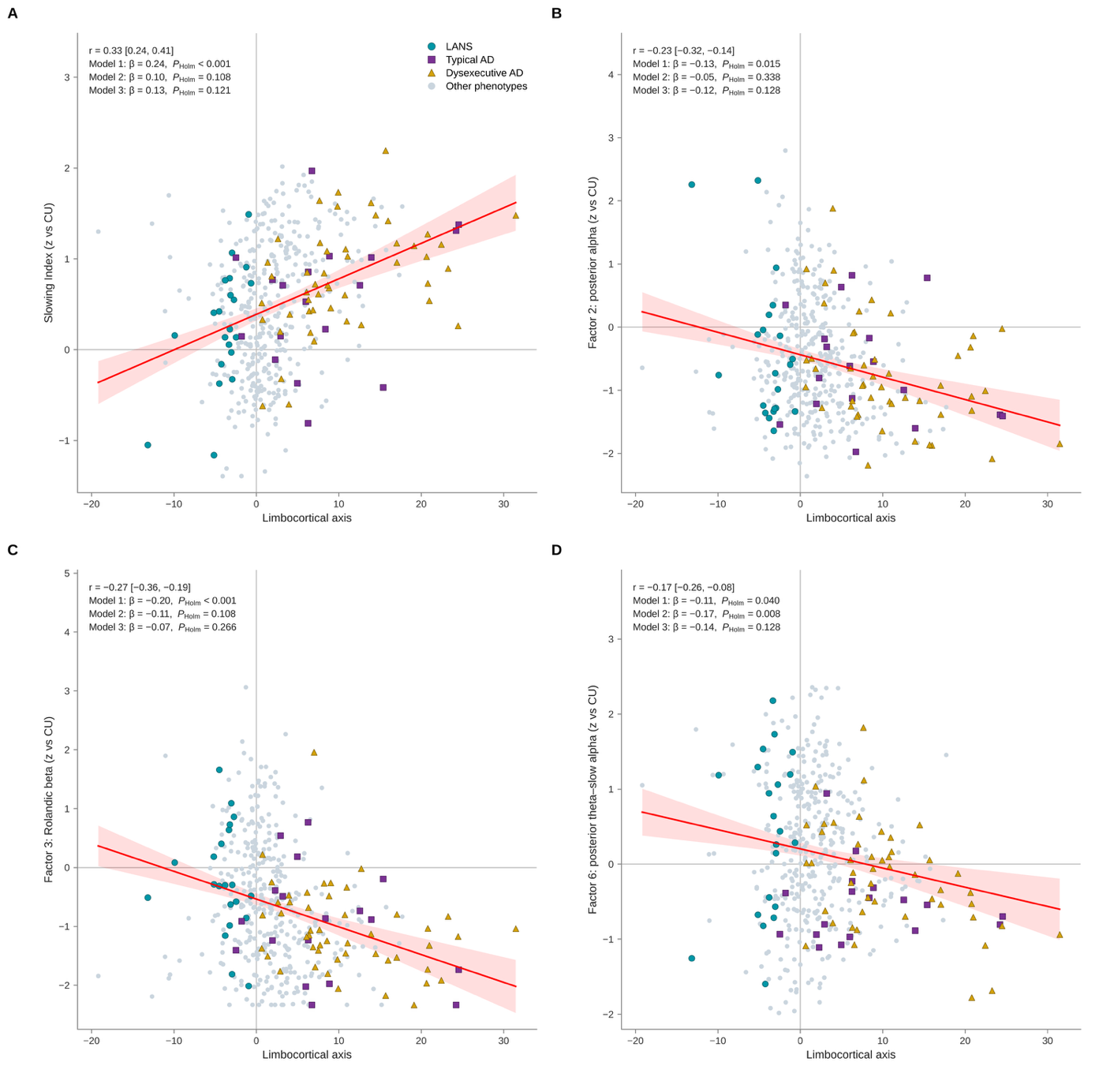

**Supplementary Figure 4: Spatial correspondence between EEG factor metabolic maps and macroscale functional gradients.** Each point is one of 400 cortical parcels (Schaefer atlas). It shows the parcel-mean β from the joint voxelwise model (factors 2, 3, 4 and 6) against the principal (G1, top row; unimodal to transmodal) or second (G2, bottom row; somatomotor to visual) gradient of Margulies et al. Lines join decile medians. Panel titles give Spearman's ρ and the two-sided spin-test *P* (10,000 rotations); an asterisk marks Holm-adjusted *P* < 0.05 across the four factors. Only factor 4 aligned with G1, with relatively higher metabolism in unimodal cortex and lower metabolism in transmodal cortex. Its V-shaped relationship with G2 is the inverse of the G1–G2 relationship itself and follows from this alignment. That correspondence was strongest on the somatomotor arm and did not survive correction.

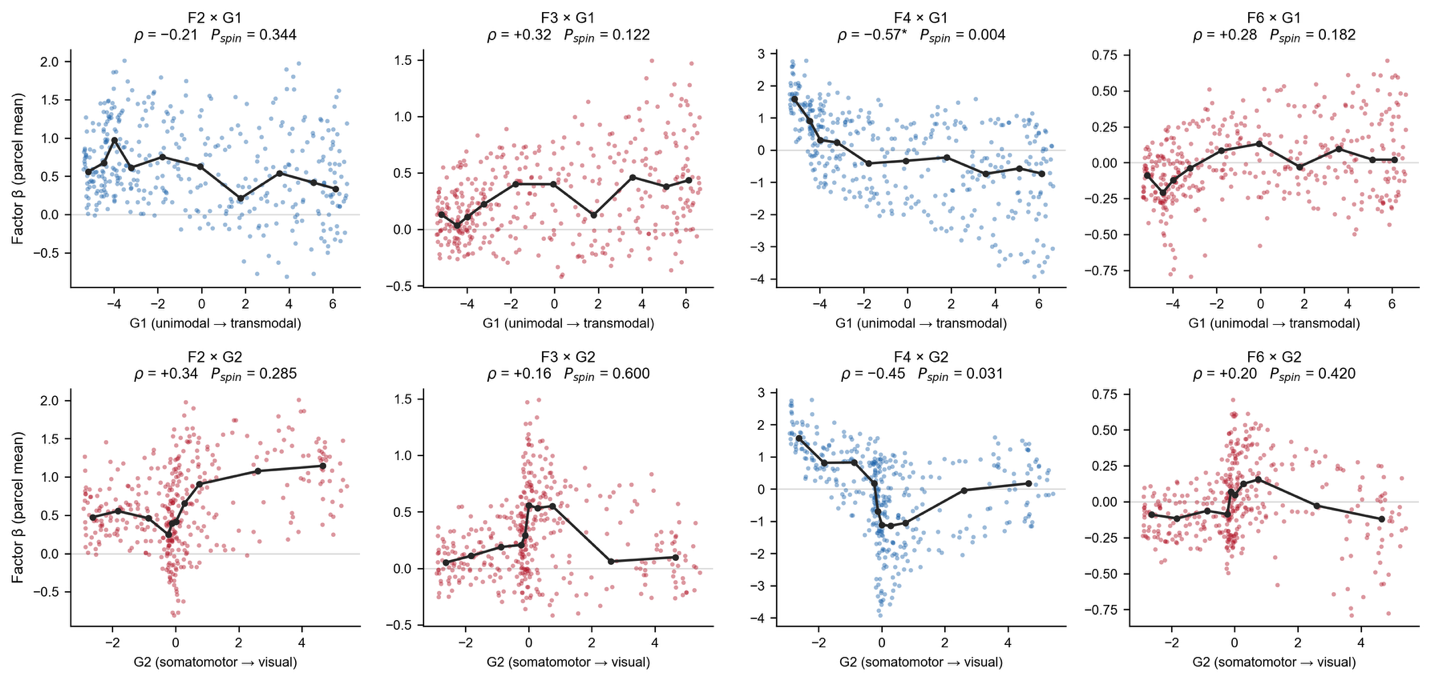

**Supplementary Figure 5: EEG factor associations with neuropsychological performance: model comparison and test-level results. (A–C)** Spearman correlations between each EEG factor and each domain composite and the global composite, shown unadjusted **(A)**, adjusted for age and education **(B)**, and additionally adjusted for FDG-PET phenotype, i.e. the within-phenotype association **(C)**. **(D)** Partial Spearman correlations (adjusted for age and education) between each factor and each individual test; the number of participants with each test is shown in parentheses. Red indicates a positive association (higher factor expression with better performance) and blue a negative association. Asterisks indicate Holm-adjusted significance within 20 comparisons per model (A–C) or 92 comparisons (D) (*p < 0.05, **p < 0.01); dots in D indicate nominal p < 0.05 that did not survive correction. AVLT, Auditory Verbal Learning Test; LM, Logical Memory; VR, Visual Reproduction; TMT, Trail Making Test; BNT, Boston Naming Test; COWAT, Controlled Oral Word Association Test; JLO, Judgment of Line Orientation; Rey–O, Rey–Osterrieth Complex Figure.

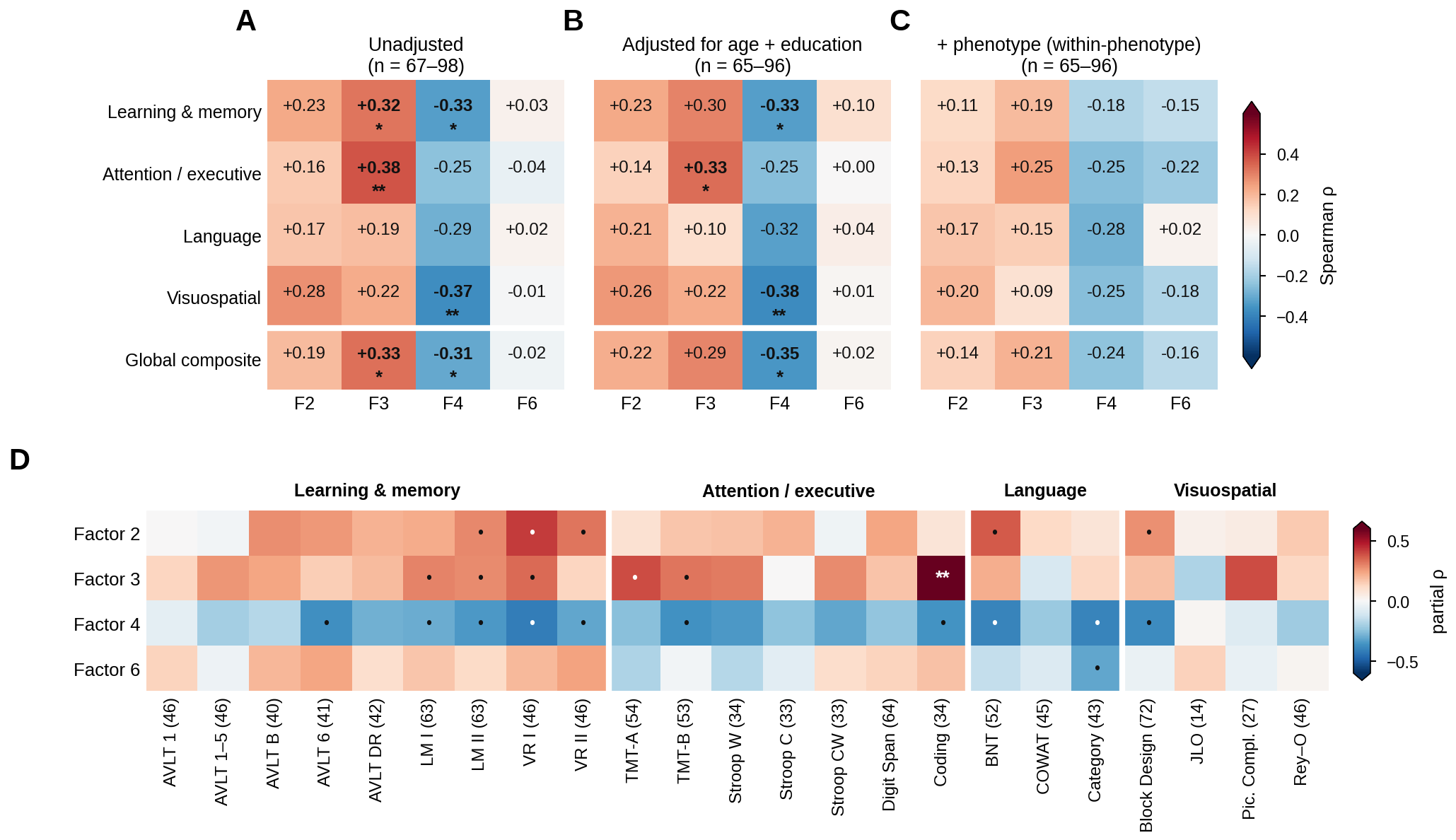

**Supplementary Figure 6: Comparison of classification algorithms.** (**A**) Cross-validated area under the receiver operating curve (AUC) for every classification analysis (rows) and classifier (columns), with the highest AUC in each row shown in bold. NB, Gaussian naïve Bayes; LR, L2-regularized logistic regression; SVM, support vector machine with radial basis function kernel; RF, random forest; GB, histogram-based gradient boosting. (**B**) Cross-validated AUC with stratified-bootstrap 95% confidence intervals for the pooled binary analyses and the four-class model (macro-averaged AUC), by classifier. Dashed lines indicate chance (AUC = 0.5). (**C**) Balanced accuracy at the class-balanced decision threshold for the same analyses. Dashed lines indicate chance: 0.5 for binary analyses and 0.25 for the four-class model.

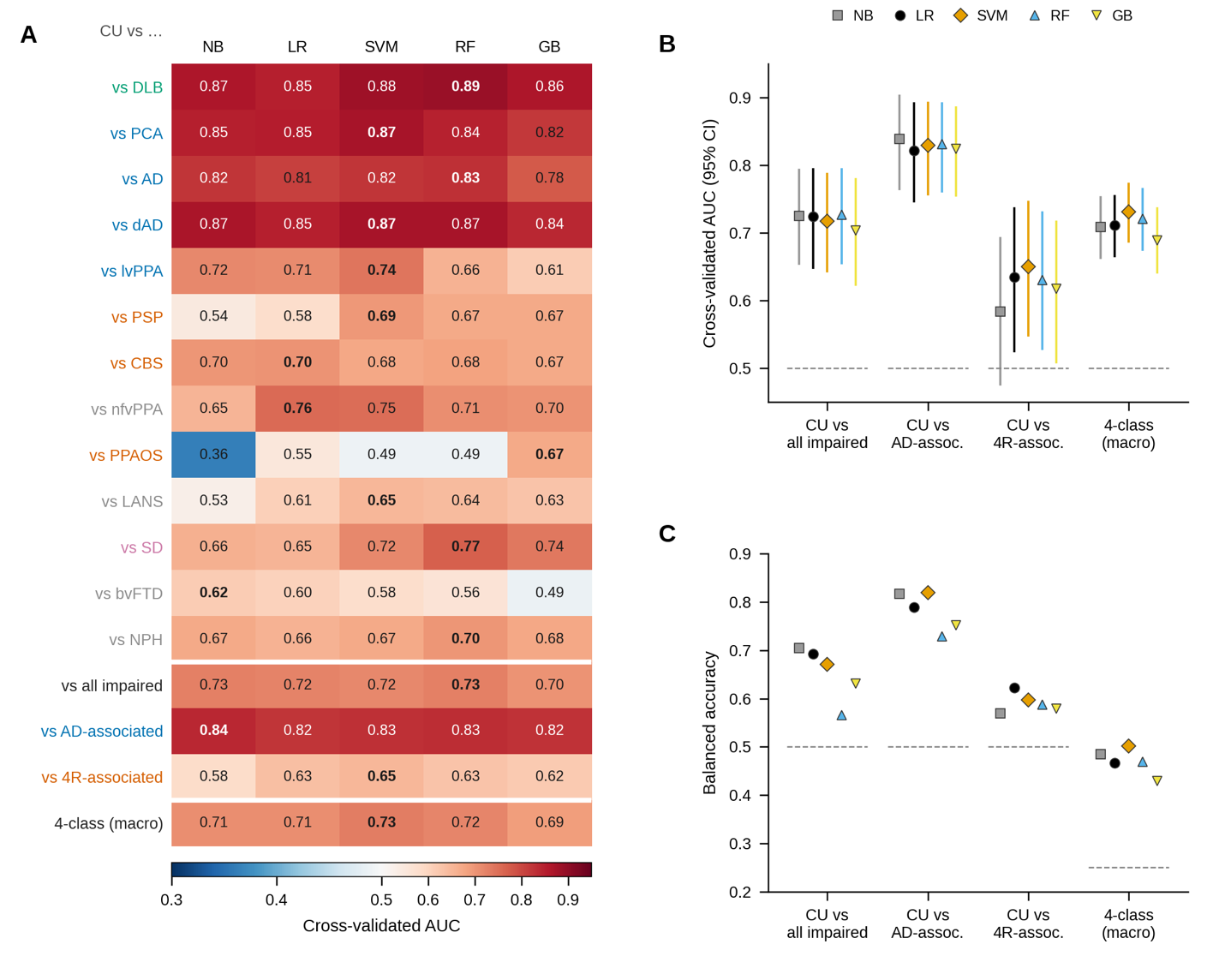

**Supplementary Tables**

**Supplementary Table 1 Number of cases retained at each p-value threshold, by group**

| **Group** | **p < 0.5** | **p < 0.1** | **p < 0.01** | **Kept** |
| --- | --- | --- | --- | --- |
| PPAOS | 99 | 50 | 17 | 17% |
| CBS | 119 | 54 | 22 | 18% |
| LANS | 80 | 72 | 21 | 26% |
| PSP | 53 | 45 | 25 | 47% |
| bvFTD | 43 | 35 | 20 | 47% |
| nfvPPA | 36 | 32 | 18 | 50% |
| SD | 55 | 49 | 37 | 67% |
| CU | 74 | 74 | 50 | 68% |
| lvPPA | 32 | 30 | 22 | 69% |
| dAD | 70 | 58 | 48 | 69% |
| NPH | 84 | 79 | 67 | 80% |
| DLB | 59 | 58 | 47 | 80% |
| PCA | 52 | 47 | 43 | 83% |
| AD | 19 | 19 | 18 | 95% |
| **Total** | **875** | **702** | **455** | **52%** |

Values are n. Kept = n at p < 0.01 as a percentage of n at p < 0.5. AD, Alzheimer disease; bvFTD, behavioral variant frontotemporal dementia; CBS, corticobasal syndrome; CU, cognitively unimpaired; dAD, dysexecutive Alzheimer disease; DLB, dementia with Lewy bodies; LANS, limbic predominant amnestic neurodegenerative syndrome; lvPPA, logopenic variant primary progressive aphasia; nfvPPA, nonfluent variant primary progressive aphasia; NPH, normal pressure hydrocephalus; PCA, posterior cortical atrophy; PPAOS, primary progressive apraxia of speech; PSP, progressive supranuclear palsy; SD, semantic dementia.

**Supplementary Table 2 Characteristics of the participants who underwent neuropsychological testing**

|  | **Neuropsychological testing** | **No neuropsychological testing** | **P-value** |
| --- | --- | --- | --- |
|  | **n = 98** | **n = 357** |  |
| **Demographics** | | |  |
| Age, mean (range) | 61.2 (19.1 – 86.1) | 62.2 (18.2 – 90.4) | 0.57 |
| Female sex, n (%) | 51 (52.0) | 169 (47.3) | 0.48 |
| Education, years, mean ± SD | 14.8 ± 2.8 (n = 96) | 14.8 ± 2.5 (n = 49) | 0.98 |
| **Race, n (%)** | | |  |
| White | 92 (93.9) | 322 (90.2) | 0.32 |
| Others | 6 (6.1) | 35 (9.8) |  |
| **Disease Label, n (%)** | | | 0.33 |
| Alzheimer disease | 4 (4.1) | 14 (3.9) |  |
| Behavioral variant frontotemporal dementia | 6 (6.1) | 14 (3.9) |  |
| Corticobasal syndrome | 5 (5.1) | 17 (4.8) |  |
| Cognitively unimpaired | 10 (10.2) | 40 (11.2) |  |
| Dysexecutive Alzheimer disease | 15 (15.3) | 33 (9.2) |  |
| Dementia with Lewy bodies | 6 (6.1) | 41 (11.5) |  |
| Limbic predominant amnestic neurodegenerative syndrome | 4 (4.1) | 17 (4.8) |  |
| Logopenic variant primary progressive aphasia | 2 (2.0) | 20 (5.6) |  |
| Nonfluent variant primary progressive aphasia | 7 (7.1) | 11 (3.1) |  |
| Normal pressure hydrocephalus | 14 (14.3) | 53 (14.8) |  |
| Posterior cortical atrophy | 11 (11.2) | 32 (9.0) |  |
| Primary progressive apraxia of speech | 1 (1.0) | 16 (4.5) |  |
| Progressive supranuclear palsy | 4 (4.1) | 21 (5.9) |  |
| Semantic dementia | 9 (9.2) | 28 (7.8) |  |
| **Interval between FDG-PET and neuropsychological testing** | | |  |
| Days, mean ± SD | 54.2 ± 70.0 | — |  |
| Days, median (IQR) | 23 (1–82) | — |  |
| Days, range | 0–271 | — |  |
| Testing before / same day / after, n | 46 / 7 / 41 | — |  |

Values are presented as mean (range), mean ± SD, or n (%). The neuropsychological subset comprises the participants with at least one neuropsychological test score linked to the paired EEG and FDG-PET studies; the remaining participants of the study population are shown for comparison.

P-values compare the two groups: Welch t-test for age and education, χ² test for sex, Fisher exact test for race, and a permutation χ² test (10 000 permutations) across the 14 disease labels.

Education was recorded for 96 and 49 participants in the two groups, respectively. The interval between FDG-PET and neuropsychological testing is the absolute number of days between the two and was available for 94 of 98 participants in the neuropsychological subset.

**Supplementary Table 3 Classification performance of all classifiers across analyses**

| **Classifier** | **AUC (95% CI)** | **AUC SD across repeats** | **Balanced accuracy** | **Sensitivity** | **Specificity** | ***P*** | ***P* (Holm)** |
| --- | --- | --- | --- | --- | --- | --- | --- |
| **Analysis 1: CU versus each phenotype** | | | | | | | |
| *CU (n = 50) vs DLB (n = 47)* | | | | | | | |
| Gaussian naïve Bayes | 0.87 (0.77–0.94) | 0.006 | 0.87 | 0.85 | 0.88 | — | — |
| Logistic regression (primary) | 0.85 (0.76–0.93) | 0.012 | 0.85 | 0.87 | 0.82 | 0.001 | 0.013 |
| SVM (RBF kernel) | 0.88 (0.80–0.95) | 0.009 | 0.85 | 0.81 | 0.90 | — | — |
| Random forest | 0.89 (0.80–0.95) | 0.010 | 0.87 | 0.83 | 0.92 | — | — |
| Gradient boosting | 0.86 (0.77–0.94) | 0.017 | 0.82 | 0.79 | 0.86 | — | — |
| *CU (n = 50) vs PCA (n = 43)* | | | | | | | |
| Gaussian naïve Bayes | 0.85 (0.76–0.93) | 0.011 | 0.84 | 0.84 | 0.84 | — | — |
| Logistic regression (primary) | 0.85 (0.76–0.93) | 0.010 | 0.84 | 0.86 | 0.82 | 0.001 | 0.013 |
| SVM (RBF kernel) | 0.87 (0.79–0.94) | 0.004 | 0.83 | 0.81 | 0.84 | — | — |
| Random forest | 0.84 (0.75–0.92) | 0.017 | 0.81 | 0.74 | 0.88 | — | — |
| Gradient boosting | 0.82 (0.72–0.90) | 0.019 | 0.78 | 0.74 | 0.82 | — | — |
| *CU (n = 50) vs AD (n = 18)* | | | | | | | |
| Gaussian naïve Bayes | 0.82 (0.67–0.94) | 0.012 | 0.79 | 0.72 | 0.86 | — | — |
| Logistic regression (primary) | 0.81 (0.67–0.92) | 0.016 | 0.72 | 0.67 | 0.78 | 0.002 | 0.018 |
| SVM (RBF kernel) | 0.82 (0.68–0.94) | 0.014 | 0.74 | 0.67 | 0.82 | — | — |
| Random forest | 0.83 (0.70–0.93) | 0.032 | 0.73 | 0.56 | 0.90 | — | — |
| Gradient boosting | 0.78 (0.63–0.91) | 0.033 | 0.73 | 0.56 | 0.90 | — | — |
| *CU (n = 50) vs dAD (n = 48)* | | | | | | | |
| Gaussian naïve Bayes | 0.87 (0.78–0.94) | 0.008 | 0.85 | 0.81 | 0.88 | — | — |
| Logistic regression (primary) | 0.85 (0.76–0.92) | 0.011 | 0.81 | 0.79 | 0.82 | 0.001 | 0.013 |
| SVM (RBF kernel) | 0.87 (0.78–0.94) | 0.005 | 0.85 | 0.81 | 0.88 | — | — |
| Random forest | 0.87 (0.79–0.93) | 0.012 | 0.81 | 0.77 | 0.84 | — | — |
| Gradient boosting | 0.84 (0.75–0.91) | 0.020 | 0.77 | 0.73 | 0.82 | — | — |
| *CU (n = 50) vs lvPPA (n = 22)* | | | | | | | |
| Gaussian naïve Bayes | 0.72 (0.57–0.86) | 0.019 | 0.73 | 0.68 | 0.78 | — | — |
| Logistic regression (primary) | 0.71 (0.57–0.85) | 0.022 | 0.70 | 0.68 | 0.72 | 0.007 | 0.049 |
| SVM (RBF kernel) | 0.74 (0.59–0.88) | 0.023 | 0.74 | 0.64 | 0.84 | — | — |
| Random forest | 0.66 (0.50–0.81) | 0.022 | 0.72 | 0.59 | 0.84 | — | — |
| Gradient boosting | 0.61 (0.45–0.78) | 0.037 | 0.68 | 0.59 | 0.76 | — | — |
| *CU (n = 50) vs PSP (n = 25)* | | | | | | | |
| Gaussian naïve Bayes | 0.54 (0.40–0.68) | 0.050 | 0.55 | 0.44 | 0.66 | — | — |
| Logistic regression (primary) | 0.58 (0.44–0.72) | 0.029 | 0.54 | 0.48 | 0.60 | 0.157 | 0.435 |
| SVM (RBF kernel) | 0.69 (0.58–0.81) | 0.036 | 0.65 | 0.64 | 0.66 | — | — |
| Random forest | 0.67 (0.54–0.79) | 0.039 | 0.57 | 0.36 | 0.78 | — | — |
| Gradient boosting | 0.67 (0.54–0.80) | 0.041 | 0.62 | 0.48 | 0.76 | — | — |
| *CU (n = 50) vs CBS (n = 22)* | | | | | | | |
| Gaussian naïve Bayes | 0.70 (0.56–0.83) | 0.029 | 0.69 | 0.59 | 0.78 | — | — |
| Logistic regression (primary) | 0.70 (0.56–0.83) | 0.024 | 0.69 | 0.64 | 0.74 | 0.012 | 0.072 |
| SVM (RBF kernel) | 0.68 (0.53–0.81) | 0.022 | 0.67 | 0.64 | 0.70 | — | — |
| Random forest | 0.68 (0.53–0.82) | 0.020 | 0.72 | 0.59 | 0.84 | — | — |
| Gradient boosting | 0.67 (0.52–0.81) | 0.026 | 0.68 | 0.59 | 0.76 | — | — |
| *CU (n = 50) vs nfvPPA (n = 18)* | | | | | | | |
| Gaussian naïve Bayes | 0.65 (0.50–0.80) | 0.032 | 0.64 | 0.56 | 0.72 | — | — |
| Logistic regression (primary) | 0.76 (0.62–0.87) | 0.021 | 0.66 | 0.61 | 0.70 | 0.001 | 0.013 |
| SVM (RBF kernel) | 0.75 (0.59–0.89) | 0.023 | 0.70 | 0.67 | 0.74 | — | — |
| Random forest | 0.71 (0.56–0.85) | 0.035 | 0.60 | 0.39 | 0.82 | — | — |
| Gradient boosting | 0.70 (0.55–0.85) | 0.042 | 0.63 | 0.44 | 0.82 | — | — |
| *CU (n = 50) vs PPAOS (n = 17)* | | | | | | | |
| Gaussian naïve Bayes | 0.36 (0.21–0.52) | 0.055 | 0.47 | 0.41 | 0.52 | — | — |
| Logistic regression (primary) | 0.55 (0.39–0.71) | 0.042 | 0.55 | 0.53 | 0.58 | 0.147 | 0.435 |
| SVM (RBF kernel) | 0.49 (0.34–0.63) | 0.068 | 0.53 | 0.47 | 0.58 | — | — |
| Random forest | 0.49 (0.33–0.65) | 0.072 | 0.51 | 0.12 | 0.90 | — | — |
| Gradient boosting | 0.67 (0.52–0.81) | 0.051 | 0.58 | 0.35 | 0.80 | — | — |
| *CU (n = 50) vs LANS (n = 21)* | | | | | | | |
| Gaussian naïve Bayes | 0.53 (0.37–0.69) | 0.042 | 0.62 | 0.62 | 0.62 | — | — |
| Logistic regression (primary) | 0.61 (0.45–0.75) | 0.030 | 0.59 | 0.57 | 0.60 | 0.145 | 0.435 |
| SVM (RBF kernel) | 0.65 (0.51–0.78) | 0.046 | 0.61 | 0.57 | 0.64 | — | — |
| Random forest | 0.64 (0.50–0.77) | 0.041 | 0.60 | 0.38 | 0.82 | — | — |
| Gradient boosting | 0.63 (0.48–0.76) | 0.046 | 0.62 | 0.43 | 0.82 | — | — |
| *CU (n = 50) vs SD (n = 37)* | | | | | | | |
| Gaussian naïve Bayes | 0.66 (0.54–0.78) | 0.023 | 0.68 | 0.68 | 0.68 | — | — |
| Logistic regression (primary) | 0.65 (0.53–0.77) | 0.021 | 0.66 | 0.68 | 0.64 | 0.023 | 0.115 |
| SVM (RBF kernel) | 0.72 (0.60–0.82) | 0.029 | 0.64 | 0.59 | 0.68 | — | — |
| Random forest | 0.77 (0.66–0.87) | 0.018 | 0.68 | 0.62 | 0.74 | — | — |
| Gradient boosting | 0.74 (0.63–0.84) | 0.023 | 0.66 | 0.62 | 0.70 | — | — |
| *CU (n = 50) vs bvFTD (n = 20)* | | | | | | | |
| Gaussian naïve Bayes | 0.62 (0.47–0.77) | 0.027 | 0.62 | 0.55 | 0.70 | — | — |
| Logistic regression (primary) | 0.60 (0.46–0.76) | 0.047 | 0.60 | 0.55 | 0.66 | 0.038 | 0.152 |
| SVM (RBF kernel) | 0.58 (0.42–0.74) | 0.051 | 0.57 | 0.50 | 0.64 | — | — |
| Random forest | 0.56 (0.41–0.73) | 0.043 | 0.59 | 0.35 | 0.84 | — | — |
| Gradient boosting | 0.49 (0.34–0.65) | 0.041 | 0.47 | 0.25 | 0.70 | — | — |
| *CU (n = 50) vs NPH (n = 67)* | | | | | | | |
| Gaussian naïve Bayes | 0.67 (0.58–0.76) | 0.014 | 0.66 | 0.57 | 0.76 | — | — |
| Logistic regression (primary) | 0.66 (0.56–0.75) | 0.010 | 0.63 | 0.61 | 0.64 | 0.005 | 0.040 |
| SVM (RBF kernel) | 0.67 (0.57–0.76) | 0.013 | 0.62 | 0.54 | 0.70 | — | — |
| Random forest | 0.70 (0.61–0.79) | 0.022 | 0.64 | 0.69 | 0.60 | — | — |
| Gradient boosting | 0.68 (0.58–0.77) | 0.025 | 0.67 | 0.67 | 0.66 | — | — |
| **Analyses 2–4: CU versus pooled phenotype groups** | | | | | | | |
| *CU (n = 50) vs all impaired (n = 405)* | | | | | | | |
| Gaussian naïve Bayes | 0.73 (0.65–0.79) | 0.010 | 0.70 | 0.65 | 0.76 | — | — |
| Logistic regression (primary) | 0.72 (0.65–0.79) | 0.012 | 0.69 | 0.69 | 0.70 | 0.001 | 0.004 |
| SVM (RBF kernel) | 0.72 (0.64–0.79) | 0.014 | 0.67 | 0.68 | 0.66 | — | — |
| Random forest | 0.73 (0.65–0.79) | 0.020 | 0.57 | 0.95 | 0.18 | — | — |
| Gradient boosting | 0.70 (0.62–0.78) | 0.016 | 0.63 | 0.86 | 0.40 | — | — |
| *CU (n = 50) vs AD-associated (n = 131)* | | | | | | | |
| Gaussian naïve Bayes | 0.84 (0.76–0.90) | 0.006 | 0.82 | 0.79 | 0.84 | — | — |
| Logistic regression (primary) | 0.82 (0.75–0.89) | 0.007 | 0.79 | 0.78 | 0.80 | 0.001 | 0.004 |
| SVM (RBF kernel) | 0.83 (0.76–0.89) | 0.009 | 0.82 | 0.78 | 0.86 | — | — |
| Random forest | 0.83 (0.76–0.89) | 0.013 | 0.73 | 0.82 | 0.64 | — | — |
| Gradient boosting | 0.82 (0.76–0.89) | 0.014 | 0.75 | 0.82 | 0.68 | — | — |
| *CU (n = 50) vs 4R-associated (n = 64)* | | | | | | | |
| Gaussian naïve Bayes | 0.58 (0.48–0.69) | 0.020 | 0.57 | 0.50 | 0.64 | — | — |
| Logistic regression (primary) | 0.63 (0.53–0.74) | 0.019 | 0.62 | 0.62 | 0.62 | 0.011 | 0.011 |
| SVM (RBF kernel) | 0.65 (0.55–0.75) | 0.020 | 0.60 | 0.59 | 0.60 | — | — |
| Random forest | 0.63 (0.53–0.73) | 0.026 | 0.59 | 0.66 | 0.52 | — | — |
| Gradient boosting | 0.62 (0.51–0.72) | 0.038 | 0.58 | 0.58 | 0.58 | — | — |
| **Analysis 5: Four-class model** | | | | | | | |
| *AD-associated (n = 131) vs 4R-associated (n = 64) vs DLB (n = 47) vs SD (n = 37)* | | | | | | | |
| Gaussian naïve Bayes | 0.71 (0.66–0.75) | 0.005 | 0.48 | — | — | — | — |
| Logistic regression (primary) | 0.71 (0.67–0.75) | 0.005 | 0.47 | — | — | 0.001 | 0.004 |
| SVM (RBF kernel) | 0.73 (0.69–0.77) | 0.009 | 0.50 | — | — | — | — |
| Random forest | 0.72 (0.68–0.76) | 0.009 | 0.47 | — | — | — | — |
| Gradient boosting | 0.69 (0.64–0.74) | 0.014 | 0.43 | — | — | — | — |

Values are cross-validated estimates from 10 × repeated stratified five-fold cross-validation, computed from each participant's out-of-fold scores averaged across repeats. AUC 95% confidence intervals were obtained by stratified bootstrap (2000 resamples). AUC SD across repeats is the standard deviation of the AUC across the 10 cross-validation repeats. Balanced accuracy, sensitivity and specificity are reported at the class-balanced decision threshold, with the patient group as the positive class. For the four-class model, AUC is the macro-averaged one-versus-rest AUC and balanced accuracy is the mean per-class recall (chance = 0.25); sensitivity and specificity are not applicable (—).

*P* = uncorrected label-permutation *P*-value (1000 permutations) for the primary model (logistic regression); *P* (Holm) = Holm-corrected within Analysis 1 (13 tests) and across Analyses 2–5 (4 tests). *P*-values were not computed for the other classifiers (—). AD-associated: PCA, lvPPA, AD and dAD. 4R-associated: PSP, CBS and PPAOS.

4R = four-repeat tauopathy; AD = Alzheimer disease; AUC = area under the receiver operating characteristic curve; bvFTD = behavioral variant frontotemporal dementia; CBS = corticobasal syndrome; CU = cognitively unimpaired; dAD = dysexecutive Alzheimer disease; DLB = dementia with Lewy bodies; LANS = limbic predominant amnestic neurodegenerative syndrome; lvPPA = logopenic variant primary progressive aphasia; nfvPPA = nonfluent variant primary progressive aphasia; NPH = normal pressure hydrocephalus; PCA = posterior cortical atrophy; PPAOS = primary progressive apraxia of speech; PSP = progressive supranuclear palsy; RBF = radial basis function; SD = semantic dementia; SVM = support vector machine.
